# Astrocyte-dependent local neurite pruning and Hox gene-mediated cell death in Beat-Va neurons

**DOI:** 10.1101/2023.12.05.570324

**Authors:** Katherine S Lehmann, Madison T Hupp, Amanda Jefferson, Ya-Chen Cheng, Amy E Sheehan, Yunsik Kang, Marc R Freeman

**Affiliations:** Vollum Institute, Oregon Health & Science University, Portland, OR; Neuroscience Graduate Program, Oregon Health & Science University, Portland, OR

## Abstract

Neuronal remodeling is extensive and mechanistically diverse across the nervous systems of complex metazoans. To explore circuit refinement mechanisms, we screened for new neuronal subtypes in the *Drosophila* nervous system that undergo remodeling early in metamorphosis. We find Beat-Va_M_ neurons elaborate a highly branched neurite network during larval stages that undergoes local neurite pruning during early metamorphosis. Surprisingly, Beat-Va_M_ neurons remodel their branches despite blockade of steroid hormone signaling and instead depend on signaling from astrocytes to activate pruning. We show Beat-Va_L_ neurons undergo steroid hormone-dependent cell death in posterior but not anterior abdominal segments. Correct activation of apoptotic cell death relies on segment-specific expression of the hox gene *Abd-B*, which is capable of activating cell death in any Beat-Va_L_ neuron. Our work provides new model cells in which to study neuronal remodeling, highlights an important role for astrocytes in activating local pruning in *Drosophila* independent of steroid signaling, and defines a Hox gene-mediated mechanism for segment-specific cell elimination.

**Summary:** Lehmann et al. characterize two new populations of neurons that undergo remodeling during *Drosophila* metamorphosis. Beat-Va_M_ neurons undergo drastic neurite pruning that is largely independent of ecdysone signaling and instead is driven by astrocytes. Beat-Va_L_ neurons undergo *Abd-B* mediated, caspase driven cell death in a segmentally restricted manner.

## Introduction

During development, the nervous system is initially populated by too many neurons that form an excessive number of synaptic connections. This neural circuitry is subsequently refined, often in response to activity-dependent signaling mechanisms, through the elimination of exuberant synapses (Stevens, Allen et al. 2007), neurites (Williams and Truman 2005, Stanfield, O’Leary et al. 1982), or entire neurons (Karcavich and Doe 2005). Importantly, while neuronal remodeling occurs in all complex metazoans and provides a mechanism for optimization of neuronal numbers and neural circuit connectivity, aberrant neuronal remodeling is believed to underlie neurological conditions like autism spectrum disorders, schizophrenia, and epilepsy (Atz, Rollins et al. 2007, Feinberg 1982, Ishizuka, Fujita et al. 2017, Neniskyte and Gross 2017, Sekar, Bialas et al. 2016, Selemon, Rajkowska et al. 1995, Selemon, Rajkowska et al. 1998, Winchester, Ohzeki et al. 2012, Dorothy, Emily et al. 2012).

How much neuronal remodeling occurs across the mammalian nervous system remains unclear, but it is thought to be extensive and has profound effects on the final wiring diagram (Luo and O’Leary 2005, Neukomm and Freeman 2014, Riccomagno and Kolodkin 2015). Axonal and synaptic connectivity for a given neuron can be radically altered through local pruning of axons or individual synaptic connections. For instance, retinal ganglion cells (RGCs) initially project axons to the dorsal lateral geniculate nucleus (dLGN) of the thalamus and form an excessive number of synapses on target cells, but later these are refined and exuberant RGC axons and synapses are eliminated (Dorothy, Emily et al. 2012, Stevens, Allen et al. 2007). Large-scale changes in circuit wiring can also be driven by selective cell death (Hutchins and Barger 1998, Oppenheim 1985). 40% of mouse GABAergic inhibitory neurons in the developing postnatal cortex are culled by a wave of caspase-driven cell death in the first 20 days of life (Southwell, Paredes et al. 2012). Many of these changes occur long after the neurons have integrated into neural circuits and fired for weeks. Precisely how specific neurons are selected for elimination remains unclear.

Glial cells help sculpt developing neural circuits through participating in the selective elimination of neurites, synapses, or entire neurons, although the mechanisms by which this happens remain poorly defined. In the dLGN, C1q is believed to opsonize synapses that are destined for removal, and C1q bound synapses are then recognized and phagocytosed by complement receptor 3 (CR3)-bearing microglia (Schafer et al., 2012) or by astrocytes through the MERTK and MEGF10 receptors (Chung et al., 2013). Disruption of C1q signaling leads to less robust RGC terminal segregation and the retention of excess functional synapses (Stevens, Allen et al. 2007), arguing that glia somehow play a role in promoting the final execution of synapse/axon elimination. Interestingly, C1q is not required in all brain areas that undergo developmental neural refinement. In the mouse barrel cortex, innervating axons are extensively refined into segregated somatosensory maps (Petersen 2019). In this context, however, refinement relies on fractalkine signaling between neurons and microglia, and C1q signaling is dispensable (Gunner, Cheadle et al. 2019). Finally, phagocytic cells can actively drive cells toward final execution of cell death. In *C. elegans*, engulfing hypodermal cells signal to partially dead cells to help push cells over the edge and fully execute apoptosis (Hoeppner, Hengartner et al. 2001, Reddien, Cameron et al. 2001). While not widely explored in the mammalian brain, similar roles for microglia have been identified in promoting the final execution of cell death in a subset of developing Purkinje neurons (Marín-Teva, Dusart et al. 2004).

*Drosophila* has served as an excellent system in which to explore the molecular basis of neuronal remodeling events *in vivo* (Lee, Lee et al. 1999, Lee, Marticke et al. 2000, Choi 2006, Watts, Hoopfer et al. 2003, Kirilly, Gu et al. 2009, Zhang, Wang et al. 2014, Williams and Truman 2005). At pupariation, a burst of the steroid hormone 20-hydroxyecdysone (ecdysone) drives activation of neuronal remodeling programs across the nervous system, most of which are executed during the first 12-18 hours of metamorphosis (Truman, Talbot et al. 1994, Schubiger, Wade et al. 1998, Schubiger, Tomita et al. 2003). Ecdysone binds to the Ecdysone Receptor (EcR) nuclear hormone receptor, and this event acts as a timing mechanism to coordinate the initiation of animal-wide metamorphic changes (Thummel 1996, Talbot, Swyryd et al. 1993, Riddiford and Truman 1993, Koelle, Talbot et al. 1991, Pinto-Teixeira, Konstantinides et al.). Cell-autonomous blockade of EcR signaling appears to inhibit all known neuronal cell death and local pruning events in the *Drosophila* nervous system (Yamaguchi and Miura 2015, Choi 2006, Schubiger, Wade et al. 1998, Marchetti and Tavosanis 2017, Hara, Hirai et al. 2013, Winbush and Weeks 2011, Kuo, Jan et al. 2005, Schubiger, Tomita et al. 2003). For instance, in the first 6-8 hours of metamorphosis in the central nervous system (CNS), Corazonin neurons in the ventral nerve cord (VNC) activate apoptotic death through EcR (Choi 2006, Tasdemir-Yilmaz and Freeman 2014) (Choi 2006, Lee, Sehgal et al. 2013, Lee, Sehgal et al. 2019, Lee, Wang et al. 2011, Wang, Lee et al. 2019). Likewise, mushroom body (MB) γ neurons, located in the brain lobes and involved in olfactory learning and memory, undergo stereotyped axonal and dendritic pruning (Heisenberg 1998) that can be blocked by inhibiting EcR (Lee, Marticke et al. 2000, Watts, Hoopfer et al. 2003). To our knowledge, neuronal cell types that undergo developmental remodeling independently of EcR have not been identified.

Like their mammalian counterparts, *Drosophila* glial cells play a crucial role in neuronal remodeling. First, activation of most remodeling events occurs by glial release of the Transforming Growth Factor-β (TGFβ) family member Myoglianin, which activates expression of EcR via TGFβ receptors on target neurons, thereby establishing their competence to prune upon receipt of steroid hormonal signals (Awasaki, Huang et al. 2011, Hakim, Yaniv et al. 2014, Wang, Lee et al. 2019, Yu, Gutman et al. 2013). After cell death or neurite/synapse degeneration has occurred, glial cells act as phagocytes to recognize and phagocytose neuronal debris through conserved signaling pathways like Draper/MEGF10 (MacDonald, Beach et al. 2006, Doherty, Logan et al. 2009, Hakim, Yaniv et al. 2014) or Fractalkine/Orion (Boulanger, Thinat et al. 2021, Ji, Wang et al. 2023, Perron, Carme et al. 2023).

A growing body of evidence implies that the signaling pathways engaged to drive neuronal remodeling are diverse and context-dependent, and we lack a deep understanding of the molecular basis of neuronal remodeling in any context (Schafer, Lehrman et al. 2012, Yaniv and Schuldiner 2016, Boulanger and Dura 2022). In this study, we sought to identify new cell types in which to explore new cellular and molecular mechanisms that regulate neuronal remodeling in the *Drosophila* pupal nervous system. We characterize two populations of neurons labeled by the *BeatVa-Gal4* driver—medial (Beat-Va_M_) and lateral (Beat-Va_L_) neurons—that undergo complete local neurite pruning or segment-specific apoptotic cell death, respectively. We show that local pruning in Beat-Va_M_ neurons can happen independently of EcR but requires signals from astrocytes, while segment-specific cell death of Beat-Va_M_ neurons is downstream of EcR and governed by spatially restricted expression of the Hox gene *Abd-B*. Our work establishes Beat-Va neurons as a new model to explore neuronal remodeling *in vivo* and identifies new mechanisms for regulation of developmental pruning and cell death in *Drosophila*.

## Results

### Beat-Va neurons undergo cell death or local pruning during metamorphosis

We sought to identify new Gal4 driver lines that would enable the characterization of neuronal remodeling in subsets of neurons in the ventral nerve cord (VNC) and brain during *Drosophila* metamorphosis. We searched ∼5,500 lines on the open source Janelia FlyLight website (Pfeiffer, Jenett et al. 2008) to identify driver lines that were active in sparse populations of neurons during the wandering 3^rd^ instar larval stage (wL3), the developmental stage that directly precedes metamorphosis (Figure 1A). From these, we selected 87 lines and validated their wL3 expression patterns by crossing each line to membrane-tethered GFP (*UAS-mCD8::GFP*) (Supplemental Table 1). We chose 28 lines to further investigate and visualized their morphologies during metamorphosis at 6 hours after puparium formation (APF) and head eversion (HE, ∼12 hrs APF), the time point by which most known cell types have completed remodeling, and 18 hrs APF (Figure 1A, Supplemental Table 2).

**Figure 1:**
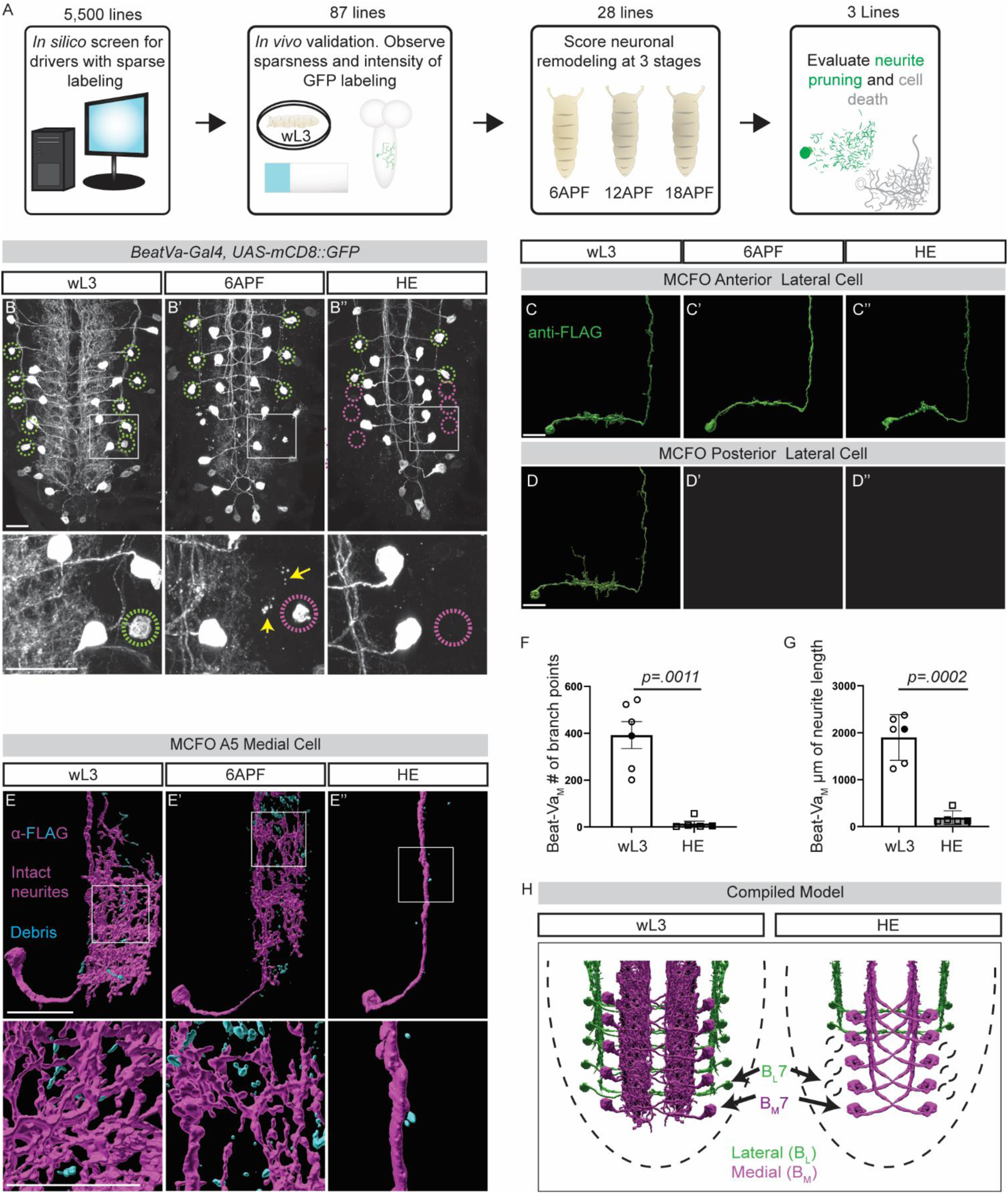
Beat-Va neurons undergo local neurite pruning or cell death. **A)** *Gal4* lines generated by Janelia were screened *in silico,* 87 drivers that label sparse populations were verified for consistency and driver strength and 28 of those were chosen for further evaluation at 6APF, 12APF (HE), and 18APF. **B)** Z-projection of ventral nerve cords labeled by *BeatVa-Gal4* driving an *UAS-mCD8::GFP* transgene at wL3 (B), 6APF (B’), and HE (B”). Surviving lateral neurons are noted by green circles, and dead or dying lateral neurons are pink. Yellow arrows denote neurite debris. **C)** Single cell morphology of the anterior Beat-Va lateral cell (Beat-Va_L_) at wL3 (C), 6APF (C’), and (C”) HE. **D)** Posterior lateral cells at wL3 (D), 6APF (D’), and HE (D”). **E)** Beat-Va medial cells (Beat-Va_M_) at wL3 (E), 6APF (E’), and HE (E”). Intact neurites in magenta and fragmented neurites in cyan. **F)** Quantification of the number of branch points in Beat-Va_M_ cells. **G)** Quantification of the total combined length of all filaments in Beat-Va_M_ cells. **H)** Composite model of both Beat-Va_L_ and Beat-Va_M_ neurons. Comparisons by student t-test Scale bar is 20 microns. Boxed region is magnified below each panel.

We focused on VNC neurons labeled by *GMR92H04-GAL4*, which was constructed by fusing an enhancer element for the *Beat-Va* gene to the DSCP and *Gal4* promoters (Jenett, Gerald et al. 2012). We refer to this line as *BeatVa-Gal4* and the neurons labeled as Beat-Va neurons. To define the segmental patterns of neurons labeled by the *BeatVa-Gal4* driver, we used antibody staining for the transcription factors Even-skipped (Eve) and Engrailed (En), which act as convenient landmarks for the identification of individual cells in the VNC (Manoukian and Krause 1992). We found that the *BeatVa-Gal4* driver labeled one lateral neuron (referred to as Beat-Va_L_) and one medial neuron (referred to as Beat-Va_M_) per hemisegment in abdominal segments A3-7 (Supplemental Figure 1A-B). This driver also weakly labeled neurons in A8 and A1-2 segments, as well as a handful of cells in the brain lobes (data not shown). The total number of cells labeled by this *BeatVa-Gal4* driver decreased prior to HE in the VNC, suggesting that a subset of neurons underwent cell death. In addition, the complexity of neurite projections in the synaptic neuropil decreased, indicating that some of these neurons underwent pruning. (Figure 1B-B’’).

To examine the morphology of these cells in segments A3-7, we used the MultiColorFlpOut (MCFO) approach. MCFO is based on the use of UAS-regulated expression of spaghetti monster GFP transgenes (smGFP) (Nern, Pfeiffer et al. 2015). Stochastic expression of one of four UAS-regulated versions of fluorescently-dead smGFP, each with a unique epitope tag (OLLAS, V5, HA, or Flag), is activated stochastically within the population of Gal4 expressing neurons (Viswanathan, Williams et al. 2015, Nern, Pfeiffer et al. 2015). Individual neurons are then visualized by staining for each epitope (Supplemental Figure 1C). In single-cell clones, we found that at 3^rd^ instar larval stages, Beat-Va_L_ cells cross the midline, project anteriorly, and terminate within the VNC. Anterior Beat-Va_L_ neurons in segments A3-4 persisted through HE (Figure 1C), while posterior lateral Beat-Va_L_ neurons in segments A5-7 underwent cell death by 6 hrs APF (Figure 1D, Supplemental Figure 1D). In contrast, we found that Beat-Va_M_ neurons extend a dense network of fine projections through multiple segments in the synaptic neuropil, totaling >1000 µm in total length. These fine processes begin fragmenting by 6 hrs APF and are cleared from the CNS by HE (Figure 1E-G).

### Local pruning in Beat-Va_M_ neurons is not driven by Ecdysone receptor (EcR)

Canonically, local neurite pruning during *Drosophila* metamorphosis depends on ecdysone signaling mediated through the Ecdysone receptor B1 (EcR-B1) (Zhu, Chen et al. 2019, Yu, Gutman et al. 2013, Kuo, Jan et al. 2005, Lee, Marticke et al. 2000). Consistent with the notion that EcR signaling also regulated Beat-Va_M_ neuron pruning, we found EcR-B1 was expressed in all Beat-Va_M_ neurons in segments A3-7 (Supplemental Figure 2A). To determine whether ecdysone signaling drove Beat-Va_M_ remodeling through EcR, we used a *UAS-EcR^DN^*construct to cell-autonomously block EcR signaling in all Beat-Va neurons. We found that expression of EcR^DN^ appeared to reduce the total number of cells eliminated in posterior abdominal segments but did not block the pruning of Beat-Va_M_ neurite fine processes (Figure 2A-B). To visualize individual cells more precisely, we generated single-cell clones in Beat-Va_M_ neurons and quantified neuronal complexity in wL3 stages and HE using the MCFO approach. At wL3 stages, we found no differences in Beat-Va_M_ neuron morphology and neurite complexity when we compared controls to EcR^DN^ expressing cells, regardless of the segment (A3-7), arguing that EcR^DN^ expression throughout larval stages does not alter Beat-Va_M_ neuron development (Figure 2C-H). Surprisingly, we also found that expression of EcR^DN^ did not block the local pruning of Beat-Va_M_ neurites by HE (Figure 2A-D), as measured by total neurite length or total number of branch points (Figure 2E-F).

**Figure 2:**
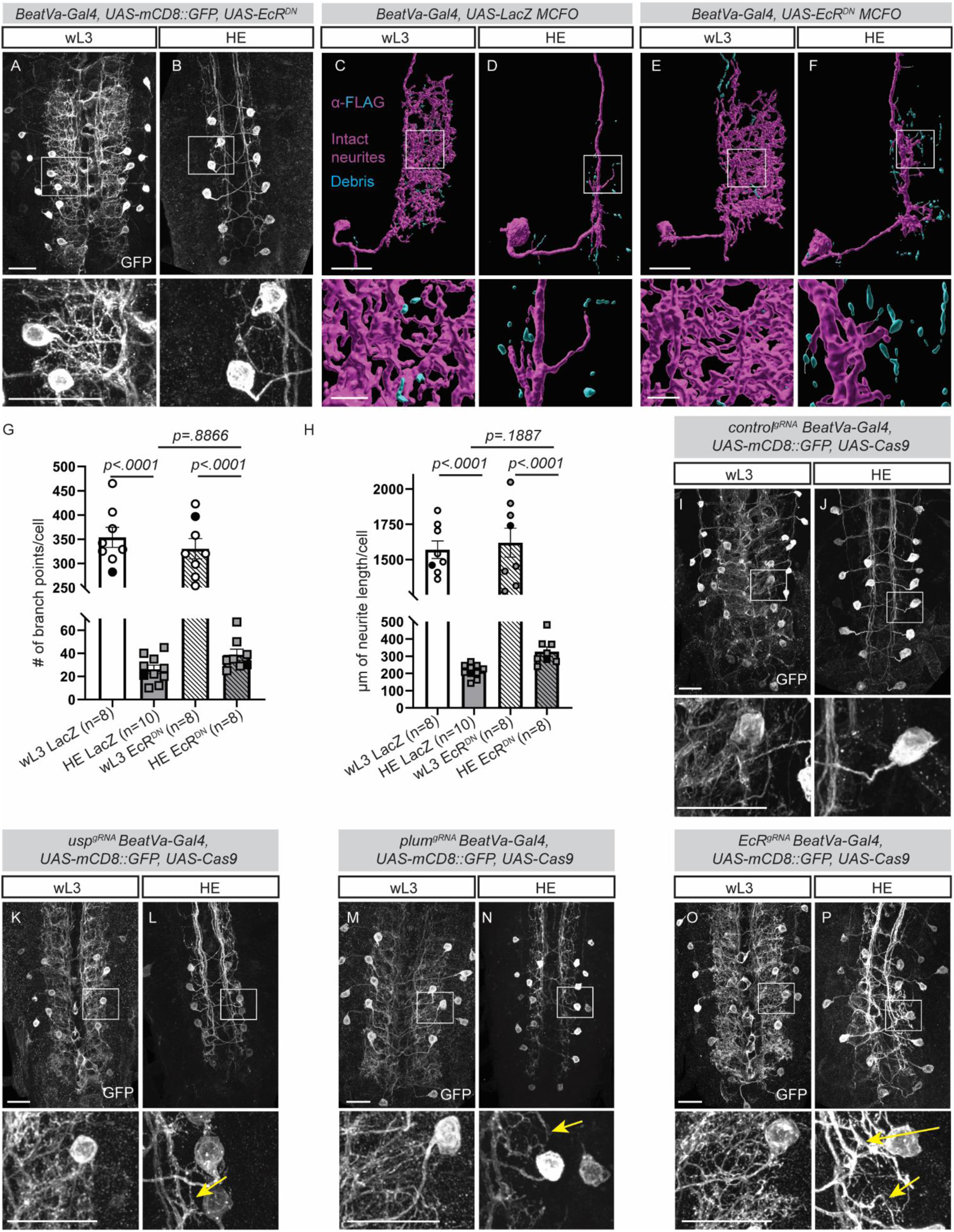
Beat-Va_M_ neurons remodel when EcR signaling or expression is inhibited. **A)** Beat-Va neurons genetically labeled with mCD8::GFP at wL3, expressing *UAS-EcR^DN^*. **B)** Beat-Va_M_ neurons expressing EcR^DN^ at HE. **C)** Control Beat-Va_M_ neuron driving *UAS-LacZ* using the MCFO technique at wL3 and HE (D). Intact neurites, magenta; fragmented neurites, cyan. Boxed area is shown in high magnification below each image. **E)** Beat-Va_M_ neuron expressing EcR^DN^ labeled with the MCFO technique at wL3 and HE (F). **G)** Quantification of Beat-Va_M_ branch point number at HE in EcR^DN^ background. **H)** Quantification of Beat-Va_M_ total neurite length in EcR^DN^ background. **I)** Beat-Va neurons genetically labeled with mCD8::GFP and expressing Cas9 under UAS control. Guide RNAs (gRNAs) are expressed ubiquitously. Cell-specific knockout of control guide RNAs at wL3 (I) and HE (J). **K)** Targeting *Ultraspiracle (usp)* with gRNAs in Beat-Va_M_ neurons at wL3 (K) or at HE (L). Fine neurites, yellow arrow. Boxed areas are displayed in high magnification below each image. **M)** *plum* gRNAs in Beat-Va_M_ neurons at wL3 (M) or HE (N). Fine neurites, yellow arrow. **O)** Expression of *EcR* gRNAs in wL3 neurons (O) and HE (P). Fine neurites, yellow arrow. Comparisons with Two-Way ANOVA and Sidak test for multiple comparisons. (A-B, I-P) Scale bars are 20 microns. (C-F) Scale bars are 20 microns in single cell images and 5 microns in the magnified view.

Given the broad role for EcR in pruning, we sought to explore the possibility that EcR^DN^ could be failing to block pruning due to insufficiently high levels of endogenous EcR expression in Beat-Va_M_ neurons. We first stained for the EcR^DN^ protein and observed robust expression of EcR^DN^ in all Beat-Va_M_ neurons at 6 hrs APF when endogenous EcR levels are low (Supplemental Figure 2B), suggesting we had sufficient EcR^DN^ expression. It was also possible that EcR^DN^ might have sufficient inherent activity to induce pruning in Beat-Va_M_ neurons (Cherbas, Hu et al. 2003), so we sought to devise alternative strategies to block EcR signaling in Beat-Va_M_ neurons. Because *EcR* mutants are lethal at late embryonic or early larval stages and because the *EcR* locus is proximal to FRT sites used for mosaic analysis, we used CRISPR/Cas9 technology to selectively eliminate genes required for EcR signaling in Beat-Va neurons (Meltzer, Marom et al. 2019). Briefly, we used lines that ubiquitously express guide RNAs (gRNAs) to *EcR*, its obligate heterodimer, *ultraspiracle* (*usp*) (Yao, Forman et al. 1993), or *plum*, an IgSF molecule that activates *EcR* transcription, and then expressed *UAS-Cas9* selectively in Beat-Va neurons with *Beat-Va-Gal4*. Targeting *EcR* and *plum* using gRNAs/Cas9 led to a significant reduction in EcR expression, as expected while targeting *usp* did not change EcR expression (Supplemental Figure 2C-D). When we examined refinement of Beat-Va_M_ neurons at HE, we observed only minor preservation or regrowth of small projections in *usp*, *EcR,* and *plum* gRNA backgrounds. (Figure 2G-N). Together, these data suggest that Beat-Va_M_ neurons activate local pruning in a manner that does not depend primarily on EcR, although the small preservation we see suggests EcR may have a minor role.

Finally, we used RNAi to concurrently knock down *BaboA* and *plum*, two transmembrane proteins that work together to induce EcR transcription, using UAS-driven RNAi constructs in *BeatVa-Gal4* animals (Figure 3A-D) (Zheng, Wang et al. 2003, Yu, Gutman et al. 2013, Wang, Lee et al. 2019). This strategy caused a strong depletion of EcR protein, as determined by antibody stains (Figure 3E). We then assessed neuronal pruning in the *BaboA/plum* double knockdown condition with MFCO and found that Beat-Va_M_ neurons continued to prune neurites (Figure 3G-K). These data, coupled with our observations using EcR^DN^ and guide RNAs/Cas9, support the notion that neurite pruning in Beat-Va_M_ neurons occurs largely independently of EcR signaling.

**Figure 3:**
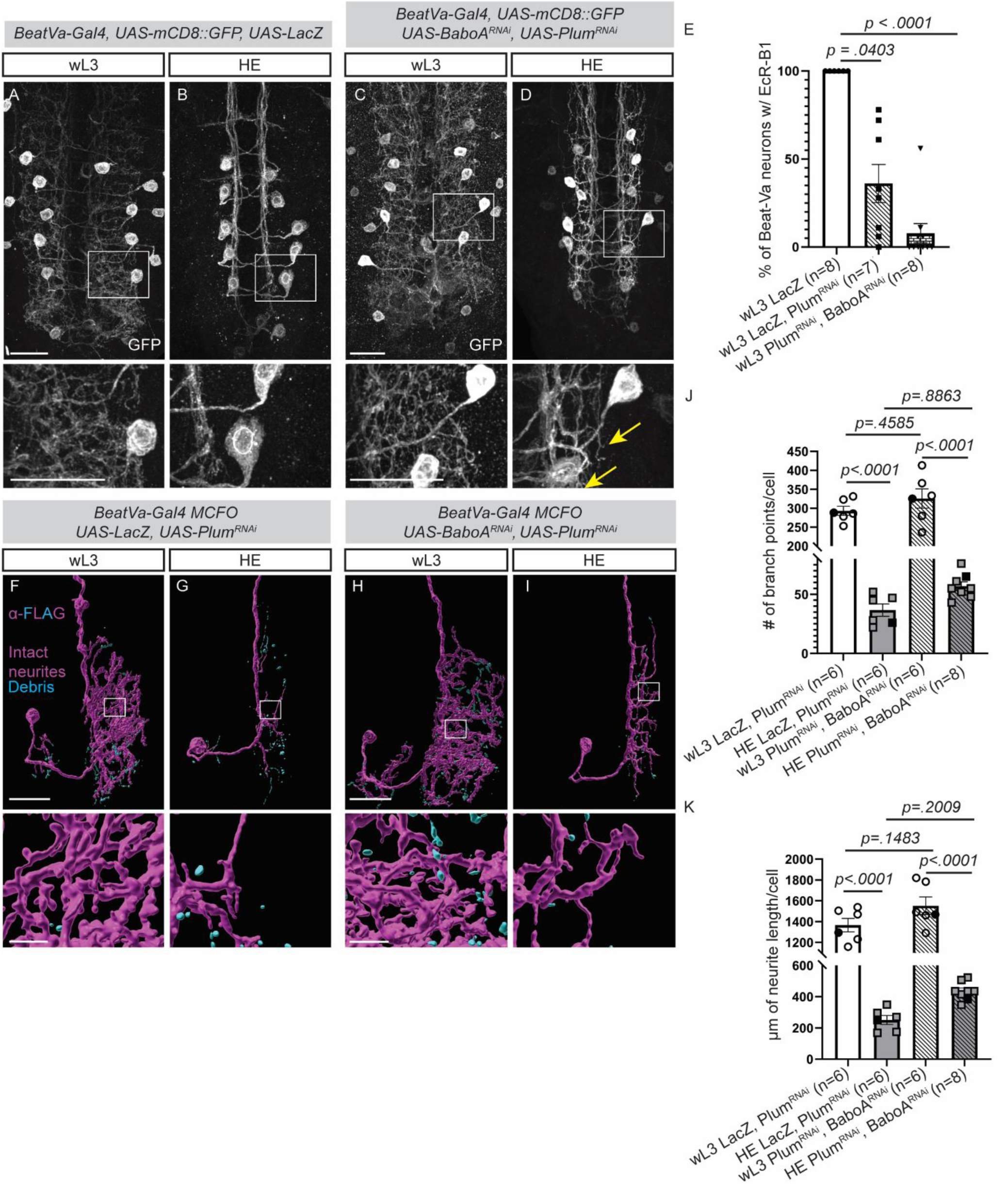
Genetic depletion of EcR by targeting upstream regulators does not block Beat-Va_M_ neuron remodeling. **A)** Beat-Va neurons labeled with mCD8::GFP at wL3 or HE (B) while driving a *UAS-LacZ* as a control construct. Boxed area shown at higher magnification below. **C)** Beat-Va neurons expressing *UAS-Plum^RNAi^* and *UAS-BaboA^RNAi^*to suppress EcR-B1 expression at wL3 and HE (D). Remaining projections, yellow arrow. Scale bars are 20 microns. **E)** Dual expression of *UAS-BaboA^RNAi^* and *UAS-Plum^RNAi^* results in substantial loss of EcR protein by antibody staining. One-way ANOVA, Kruskal-Wallis test for multiple comparisons. **F)** Beat-Va_M_ neurons driving *UAS-LacZ* and *UAS-Plum^RNAi^* labeled using MCFO at wL3 (F) and HE (G). **H)** Beat-Va_M_ neurons driving *UAS-BaboA^RNAi^*and *UAS-Plum^RNAi^* labeled using MCFO at wL3 (H) and HE (I). Intact neurites, magenta; fragmented neurites, cyan. Boxed areas are shown in high magnification below the image. **J)** Quantification of branch point number and (K) neurite length for (F-I). Comparison with Two-Way ANOVA and Sidak test for multiple comparisons. (A-D) Scale bars are 20 microns. (F-I) Scale bars are 20 microns in single cell images and 5 microns in the magnified view.

### Astrocytes non-cell-autonomously regulate Beat-Va_M_ neuron pruning

Beat-Va_M_ neurite fragmentation and clearance occurs coordinately with the EcR-dependent transformation of larval astrocytes into phagocytic glia (Tasdemir-Yilmaz and Freeman 2014, Kang 2023). To determine what role astrocytes might play in driving Beat-Va_M_ neuron remodeling, we generated a *LexAop-EcR^DN^* line, which we could drive with *alrm-LexA* to selectively block the transformation of astrocytes into phagocytes while allowing all other EcR-mediated neuronal signaling events to proceed normally. We first confirmed that our *LexA/LexAop* constructs efficiently blocked astrocyte transformation into phagocytes at 6 hrs APF (Supplemental Figure 3G-J). Indeed, expression of this EcR^DN^ construct in astrocytes resulted in astrocytes retaining their bushy morphology characteristic of larval astrocytes (Supplemental Figure 3A-F) (Hakim, Yaniv et al. 2014, Tasdemir-Yilmaz and Freeman 2014). We then used the MCFO approach to examine Beat-Va_M_ neuron architecture at wL3 and HE after blocking EcR-mediated signaling selectively in astrocytes. Strikingly, we found that blockade of EcR signaling in astrocytes strongly suppressed local pruning in Beat-Va_M_ neurons; in the absence of astrocytic EcR signaling, we found significantly less neurite remodeling in Beat-Va_M_ neurons at HE (Figure 4A-F).

**Figure 4:**
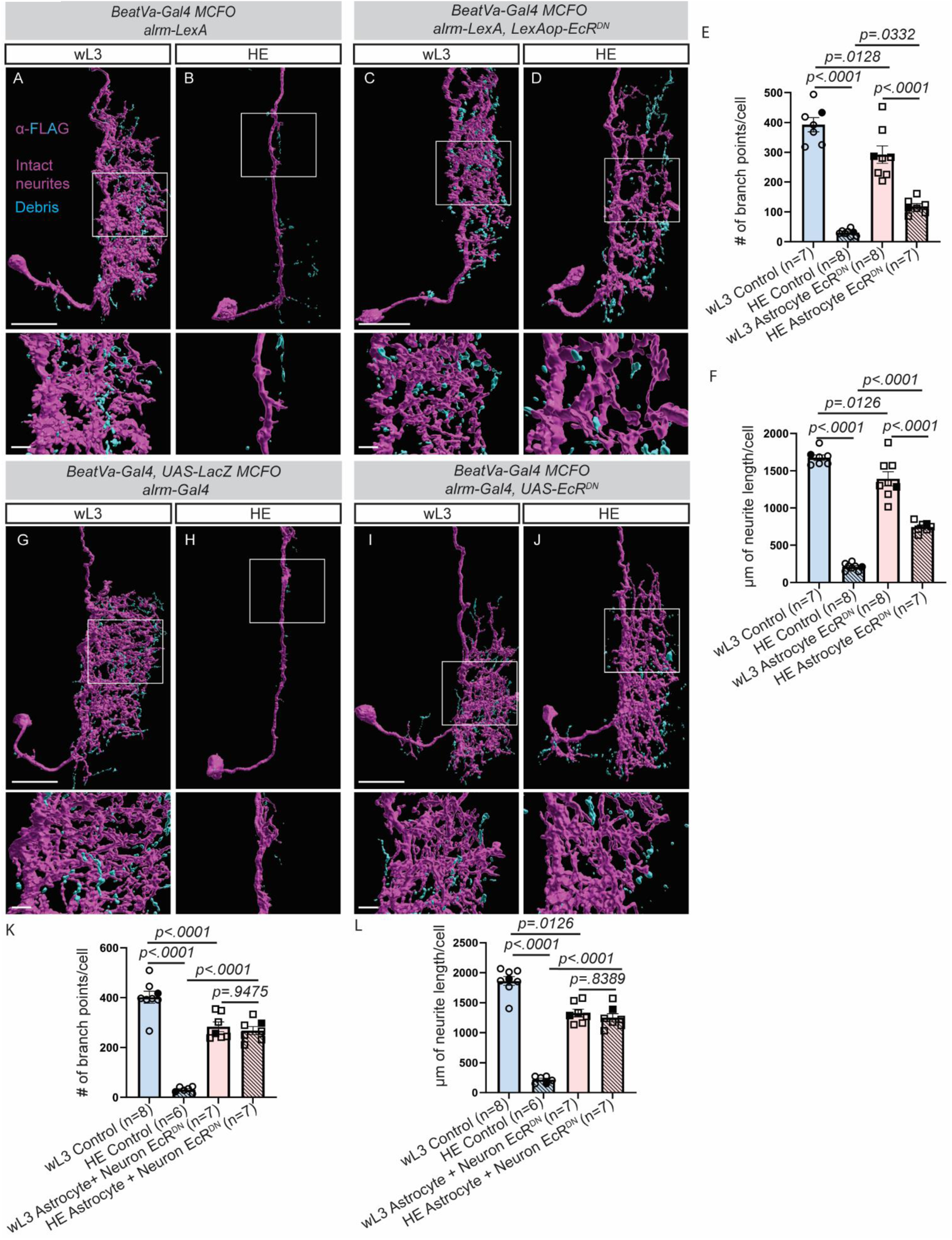
Astrocyte-derived signals converge with intrinsic Beat-Va_M_ neuron EcR signaling to execute local pruning. **A)** Beat-Va_M_ neurons labeled with MCFO in an *alrm-LexA* background (control) at wL3 (A) or HE (B). **C)** *alrm-LexA, LexAOp-EcR^DN^* background labeled with the MCFO technique at wL3 (C) or HE (D). **E)** Quantification of point number (E) or total neurite length (F) of (A-D) **G)** Beat-Va_M_ neurons in an *alrm-Gal4* background (control) at wL3 (G) HE (H). **I)** Beat-Va_M_ neurons in an *alrm-Gal4, UAS-EcR^DN^* background at wL3 (I) or HE (J). **Q)** Quantification of point number (Q) or total neurite length (R) of (G-J). All comparisons are done with Two-way ANOVA with the Sidak test for multiple comparisons. Intact neurites, magenta; fragmented neurites, cyan. Scale bars are 20 microns for whole neuron images and 5 microns for magnified images.

Given that inhibition of EcR signaling in Beat-Va_M_ neurons had a minor effect on local pruning, we reasoned that remodeling of Beat-Va_M_ neurons might be driven by a combination of EcR signaling in Beat-Va_M_ neurons and EcR-dependent signaling from astrocytes. To test this directly, we blocked both astrocyte and neuronal ecdysone signaling by driving a *UAS-EcR^DN^* simultaneously in both astrocytes and Beat-Va neurons (using *alrm-Gal4* and *BeatVa-Gal4*). We found that the combination of blocking astrocyte transformation and blocking neuronal ecdysone signaling led to a complete inhibition of local pruning in Beat-Va_M_ neurons (Figure 4G-L). Together, these data indicate that local Beat-Va_M_ neurite pruning occurs in response to EcR-dependent signaling in both Beat-Va_M_ neurons and astrocytes, with astrocytic EcR signaling playing the primary role. Finally, we note that at wL3 stages before local pruning of Beat-Va_M_ neurons, neurites in animals expressing astrocytic EcR^DN^ were simplified to a small degree in total neurite length and number of branch points compared to controls (Figure 4E-F, K-L). This observation argues that EcR-dependent signaling in astrocytes is important for a small fraction of Beat-Va_M_ neurite growth during embryonic or larval stages.

### The TGFβ molecule Myoglianin activates EcR expression in astrocytes to drive Beat-Va_M_ neuron local pruning

In previously studied *Drosophila* neurons, glial release of the TGFβ molecule Myoglianin (Myo), which signals through the two TGFβ receptors BaboA and Plum, results in upregulation of EcR in neurons so they are competent to remodel in response to ecdysone (Yu, Gutman et al. 2013, Awasaki, Huang et al. 2011). Accordingly, RNAi knockdown of Myo in all glia using *repo-Gal4* suppresses remodeling of these neurons in the *Drosophila* CNS (Awasaki, Huang et al. 2011). To explore the possibility that astrocytic Myo might be the factor regulating Beat-Va_M_ neuron pruning, we first generated a *BeatVa-LexA* line, which we used to drive *LexAop-Jupiter.sfGFP* (a line designed to visualize neuronal processes by labeling microtubules with GFP (Poe, Tang et al. 2017), to visualize Beat-Va neurons in a background where we could drive *UAS-Myo^RNAi^* in glia. We confirmed the expression pattern of the *BeatVa-LexA* line by comparing its overlap with *BeatVa-Gal4, UAS-mCD8::GFP* (Supplemental Figure 4). When we drove expression of *Myo^RNAi^* in all glia, we observed a strong suppression of Beat-Va_M_ neuron local pruning compared to controls (Figure 5A-D). However, when we drove *Myo^RNAi^*only in astrocytes, it did not affect Beat-Va_M_ neuron local pruning (Figure 5I). This could indicate that Myo is supplied to Beat-Va_M_ neurons by non-astrocytic glial subtypes, but this would not explain why EcR^DN^ expression in astrocytes alone could potently block Beat-Va_M_ neuron local pruning if Myo was a key factor. Surprisingly, when we examined astrocyte morphology in a background where *Myo^RNAi^* was expressed in all glia, we found astrocytes failed to transform but did so normally if we drove *Myo^RNAi^* only in astrocytes (Figure 5F-K). Furthermore, we found that EcR-B1 staining was absent from astrocytes when *Myo^RNAi^* was driven in all glia but normal when driven in astrocytes alone (Figure 5L-O). We interpret these data to mean that non-astrocytic glial subtypes provide astrocytes with Myo to activate EcR expression, which then drives astrocyte transformation into phagocytes and enables astrocyte regulation of Beat-Va_M_ neuron local pruning. Given that Myo knockdown in astrocytes alone did not block Beat-Va_M_ neuron local pruning, other astrocyte-derived factors likely regulate activation of Beat-Va_M_ neuron local pruning.

**Figure 5:**
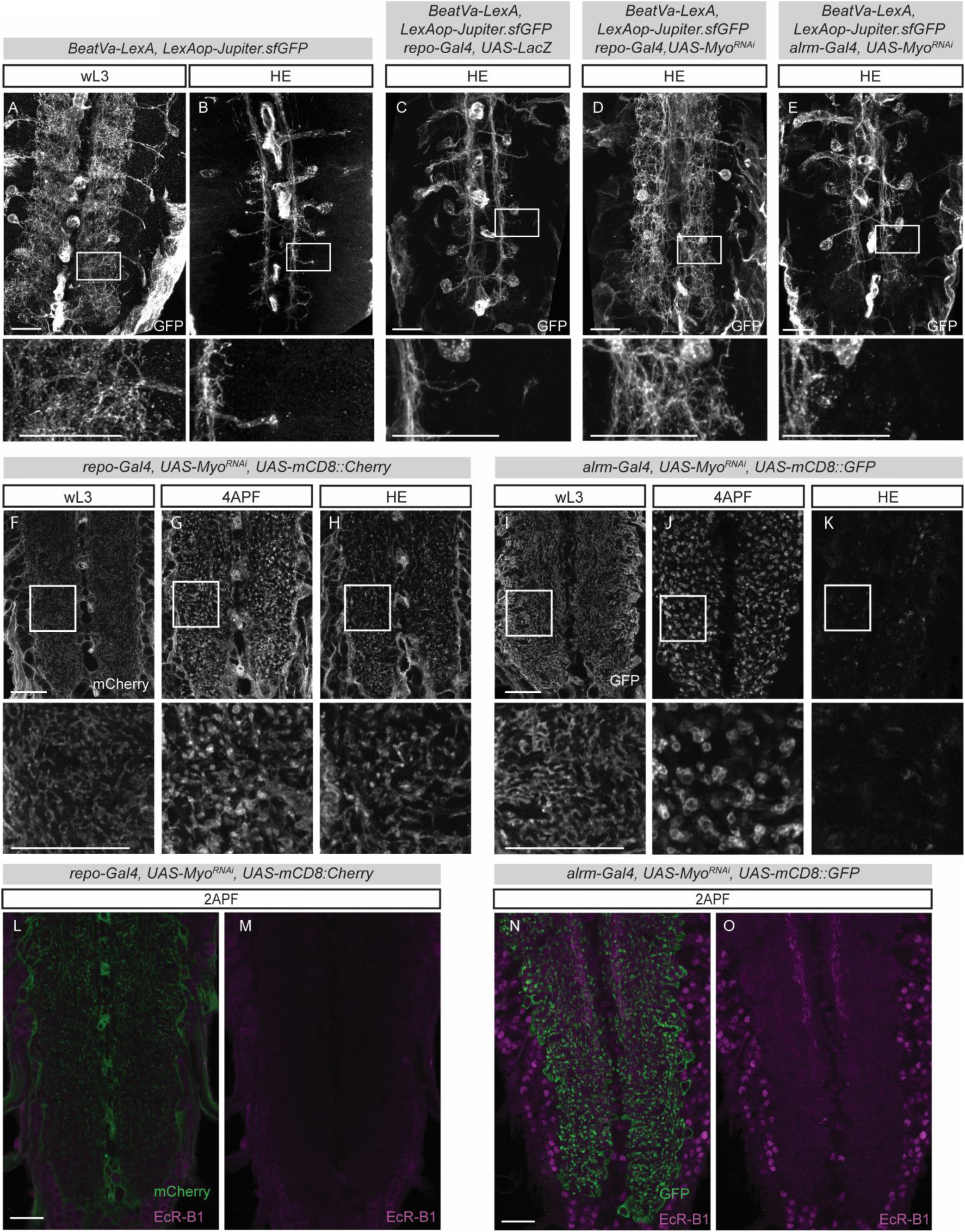
Pan-glial Myoglianin expression drives phagocytic astrocyte transformation. **A)** Beat-Va neurons labeled with a LexAop driven, microtubule targeted, superfolder GFP (*LexAop-Jupiter.sfGFP)* at wL3 or HE (B). **C)** Beat-Va_M_ neurons at HE with *repo-Gal4* and *UAS-LacZ* (control) in the background. **D)** Myoglianin knocked down in all glia with *repo-Gal4*. **E)** Myoglianin knocked down only in astrocytes with *alrm-Gal4*. F) Visualization of astrocytes using *a UAS-mCD8::Cherry* when Myoglianin is knocked down in all glia at wL3, 4 APF (G), and HE (H). **I)** Astrocytic knockdown of Myoglianin only in astrocytes at wL3, 4 APF (J) or HE (K). Note that by HE astrocytes are also no longer visible in controls. **L, M)** Expression of *UAS-Myo^RNAi^* in all glia, EcR-B1 staining, magenta. Astrocyte membranes, green. **N, O)** Expression of *UAS-Myo^RNAi^* in astrocytes, EcR-B1 staining, magenta. Astrocyte membranes, green. All scale bars are 20um.

### Beat-Va_L_ neurons are eliminated through segment-specific, steroid-dependent apoptotic cell death

Blocking ecdysone signaling with cell-specific expression of EcR^DN^ suppressed Beat-Va_L_ apoptosis (Figure 6A-C, E), similar to most other populations of *Drosophila* neurons that are eliminated by cell death during early metamorphosis (Lee, Wang et al. 2011, Winbush and Weeks, Denton, Shravage et al. 2010, Zirin, Cheng et al. 2013). To determine whether this cell death occurred by canonical apoptotic signaling mechanisms, we stained for activated caspases using an antibody that recognized a cleaved version of Dcp-1 (Peterson, Barkett et al. 2003). Posterior Beat-Va_L_ cells (A5-7) became positive for activated caspases during early metamorphosis, while cells in anterior segments that survived (A3-4) were not caspase positive (Figure 6F-H). We then expressed P35, a baculovirus caspase inhibitor that blocks many caspase-dependent cell death events (Clem, Fechheimer et al. 1991) using *BeatVa-Gal4*. We found that P35 strongly blocked cell death through HE in segments A5-6 and more weakly blocked death in A7 (Figure 6D-E) (likely due to driver strength variability), arguing posterior Beat-Va_L_ apoptosis is driven through caspase activation (Figure 6D-F, Figure 6H).

**Figure 6:**
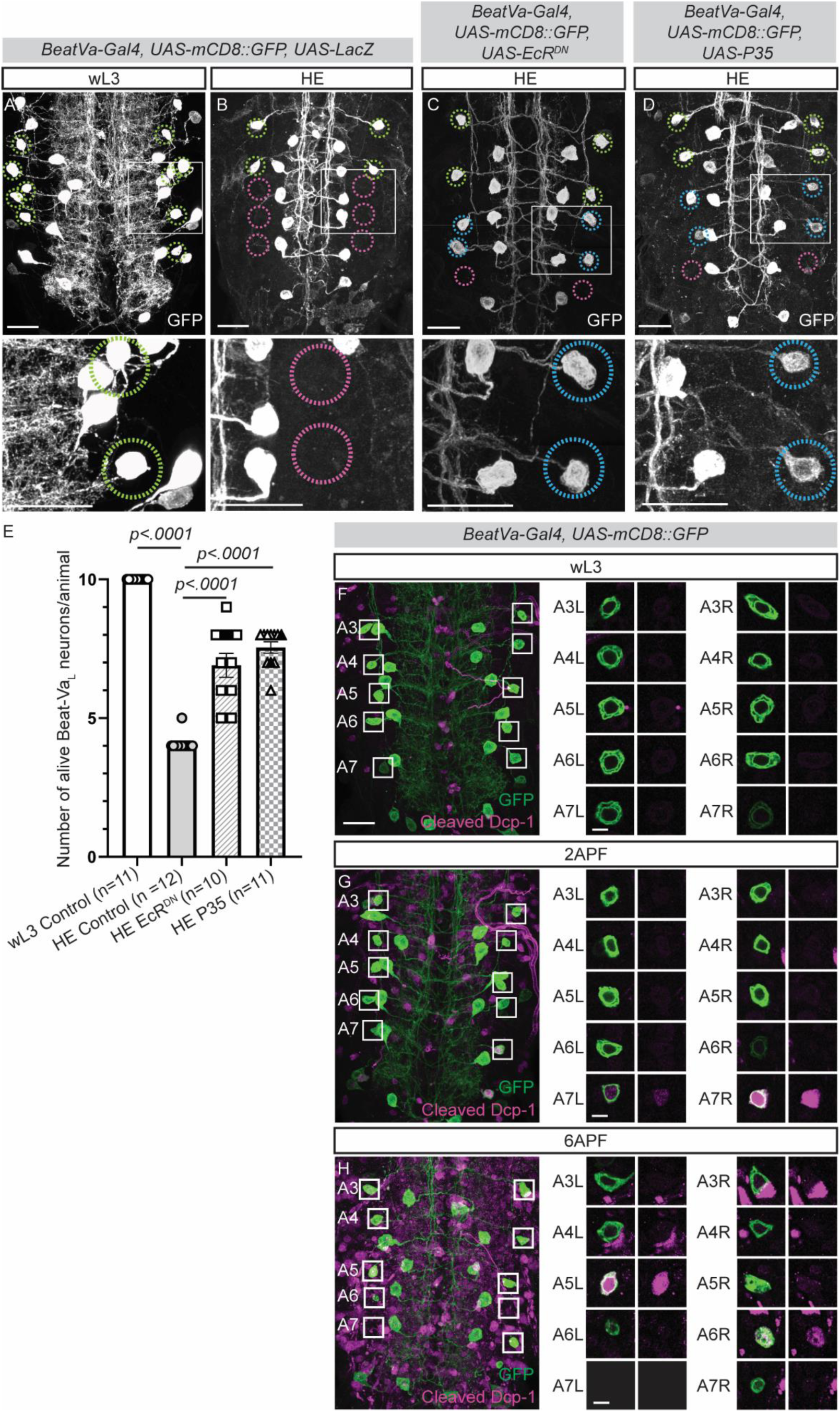
Beat-Va_L_ neurons undergo hormone-dependent, caspase-activated apoptosis. **(A)** Beat-Va neurons at wL3 (A), and HE (B) labeled with mCD8::GFP, expressing *UAS-LacZ* (control). **(C)** Beat-Va neurons at HE expressing mCD8::GFP with EcR^DN^ (C) or *UAS-P35* (D). Lime green circles indicate normal lateral cells before remodeling. Pink circles indicate the position of dead lateral cells. Blue circles indicate lateral cells surviving beyond normal cell death time point. Scale bars, 20 microns. **E)** Quantification of the number of lateral cells at wL3, or HE in controls or animal expressing EcR^DN^ or P35. Two-way ANOVA, Sidak multiple comparison test. **F)** Beat-Va neurons genetically labeled with mCD8::GFP and stained for cleaved Dcp-1 at wL3 (F), 2 APF (G) or 6 APF (H). Right, A3-A7 lateral cells from each hemisegment (L = left and R = right hemisegment) are magnified and shown as a single plane image on the right of each full VNC image. (A-D) Scale bars are 20 microns. (F-H) Scale bars are 20 microns in population images and 5 microns in the magnified view.

### Abdominal-B regulates segment-specific cell death in Beat-Va_L_ neurons

By morphological criteria and common expression of EcR-B1, Beat-Va_L_ neurons in segments A3-7 appeared similar. We speculated that positional identity and differences in survival could be regulated by differential expression of homeobox (Hox) genes. The boundary between the anterior and posterior lateral cells (A4/5) is defined by the *Abdominal-B* (A*bd-B*) Hox gene in developing embryos (Delorenzi and Bienz 1990), and during embryonic development, Abd-B can regulate apoptotic cell death of neuronal progenitors (Bakshi, Sipani et al. 2020, Clarembaux-Badell, Baladrón-de-Juan et al. 2022, Monedero Cobeta, Salmani et al. 2017). To explore the role of Abd-B in cell death of A5-7 neurons, we used antibodies to determine its larval expression pattern. We found that surviving A3-4 Beat-Va_L_ cells did not express Abd-B, while Beat-Va_L_ cells in segments A5-7 strongly expressed Abd-B at wL3 until Beat-Va_L_ cell death (Figure 7A, Supplemental Figure 5A). We then knocked down Abd-B expression specifically in the Beat-Va neurons by driving a *UAS-Abd-B^RNAi^* construct with *BeatVa-Gal4*. This led to a strong suppression of caspase activation in posterior Beat-Va_L_ neurons (detected by cleaved Dcp-1 staining) through 6 hrs APF (Figure 7B-D) and partially suppressed lateral cell death compared with controls at HE (Figure 7E-G, Figure 7I). This incomplete but strong blockade of cell death is likely due to a partial knockdown effect by the RNAi construct targeting *Abd-B,* as staining for Abd-B revealed partial rather than complete knockdown of protein levels in some cells. We note that Beat-Va_L_ cells retained expression of EcR-B1 even when Abd-B was depleted from these cells (Supplemental Figure 5B), arguing that the suppression of cell death could not be explained by changes in EcR. Finally, when we performed the reciprocal experiment and expressed Abd-B in all Beat-Va_L_ neurons by driving *UAS-Abd-B* with *BeatVa-Gal4* (Supplemental Figure 5C), we found that A3-A4 lateral cells underwent cell death by HE (Figure 7H, 7I), indicating that expression of Abd-B in Beat-Va_L_ neurons is sufficient to induce cell death during metamorphosis.

**Figure 7:**
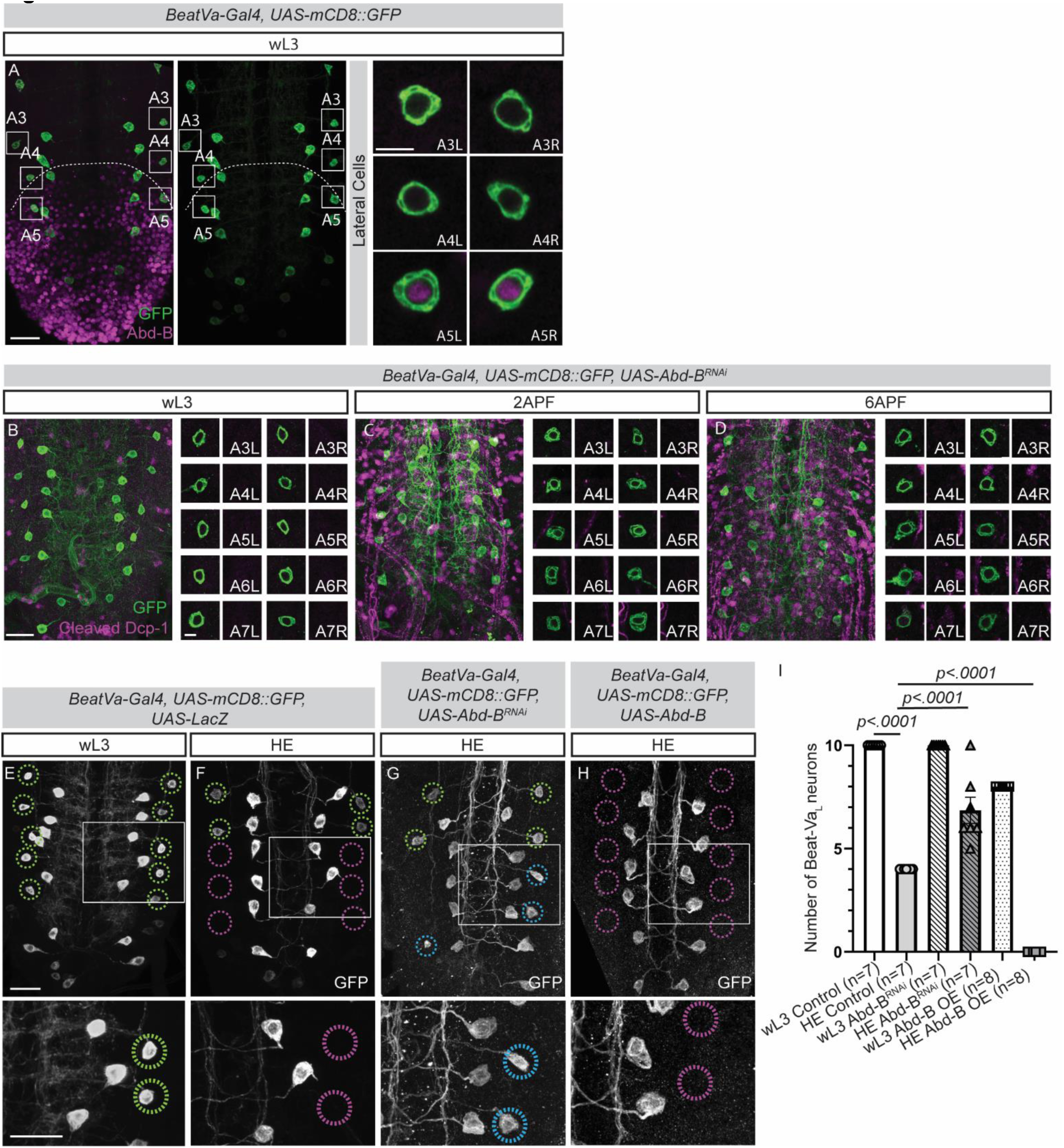
Hox gene *Abd-B* dictates caspase-dependent Beat-Va_L_ neuron cell death. **A)** Beat-Va neurons at wL3 labeled with mCD8::GFP stained with Abdominal-B antibodies (magenta). White dashed line denotes the boundary of Abdominal B (Abd-B) expression. **B)** Beat-Va neurons genetically labeled with mCD8::GFP and driving a *UAS-Abd-B^RNAi^* at wL3 (B), 2 APF (C), and 6APF (D) stained for cleaved Dcp-1. A3-A7 lateral cells from the right and left are blown up and shown as a single plane image on the right of each image. **E)** Beat-Va neurons at wL3 (E) or HE (F) genetically labeled with mCD8::GFP crossed to *UAS-LacZ* as a control. Green circles indicate Beat-Va_L_ neurons. Magenta, position of dead cells. **G)** Expression of *UAS-Abd-B^RNAi^* in Beat-Va neurons. Blue circles, cells that survive inappropriately. **H)** Expression of Abd-B in all lateral cells by HE leads to cell death. **I)** Quantification of (E-H). Two-way ANOVA, Sidak test for multiple comparisons.

(A-D) Scale bars are 20 microns in population images and 5 microns in the magnified view.

(E-H) Scale bars are 20 microns.

Beat-Va_M_ neurons in segments A5-7 also express Abd-B at wL3 and HE, although these cells do not undergo cell death (Supplemental Figure 5A, D). Furthermore, expression of Abd-B in the Beat-Va_M_ A3-4 cells (which typically do not express the protein) does not affect cell survival (Figure 7H, Supplemental Figure 5C). Finally, we observed no obvious changes in Beat-Va_M_ neurite pruning when Abd-B expression was manipulated, suggesting that Abd-B regulates cell death in some but not all populations of neurons during metamorphosis (Figure 7E-H).

## Discussion

A growing collection of mechanistic studies on neuronal remodeling suggests a diversity of molecular pathways are deployed across the nervous system to accomplish remodeling in different contexts (Schafer, Lehrman et al. 2012, Yaniv and Schuldiner 2016, Neukomm and Freeman 2014, Boulanger and Dura 2022). The relatively small number of neuronal populations in which remodeling has been studied, compared to the total number that likely exhibit remodeling across the CNS of complex metazoans, may have limited the depth of our knowledge about the diversity of signaling pathways that can drive cell death, and neurite or synaptic pruning. In this study, we identified new cell types in which we could explore the cellular phenotypes and mechanistic basis of neuronal remodeling in *Drosophila*. We described two new populations of *BeatVa-Gal4-*expressing neurons that remodel using mechanisms not previously reported in other heavily studied CNS cell types: mushroom body neurons (Lee, Marticke et al. 2000, Yu, Gutman et al. 2013, Lai, Chu et al. 2016) and ventral Corazonin neurons (Choi 2006, Lee, Sehgal et al. 2013, Wang, Lee et al. 2019). Beat-Va_M_ neurons exhibit EcR-independent, astrocyte activated local neurite pruning, while Beat-Va_L_ neurons die in a hox gene-mediated, segmentally restricted pattern.

Beat-Va_M_ neurons undergo massive local neurite pruning during the first twelve hours of metamorphosis, which we visualized with single-cell precision. These neurons are now an excellent model cell type in which to unravel the cellular and molecular mechanisms of local neurite pruning *in vivo*. Surprisingly, we found that Beat-Va_M_ neuron local pruning was not suppressed by cell-autonomous blockade of EcR signaling. However, we found that blocking astrocyte phagocytic transformation using EcR^DN^ reduced Beat-Va_M_ remodeling by ∼50%, and when combined with simultaneous blockade of EcR signaling in Beat-Va_M_ neurons, local neurite pruning was almost entirely blocked. We interpret these data to mean that extrinsic cues from astrocytes and neuron intrinsic EcR-mediated events converge to control Beat-Va_M_ remodeling.

Why a role for neuron-autonomous EcR signaling is more fully revealed when astrocytes are also blocked from transformation is an open question. A recent study showed that neuronal ecdysone signaling allows ddaC neurons to become competent for pruning by driving microtubule rearrangement, with subsequent physical force caused by tissue movement in the body wall during metamorphosis driving the severing and fragmentation of dendrites (Krämer, Wolterhoff et al. 2023). It is possible that neuronal ecdysone signaling may make Beat-Va_M_ neurons competent for remodeling through similar mechanisms, but execution of fragmentation requires a secondary mechanical event. Astrocytes could, for example, provide the physical force that severs small neurites during metamorphosis as they transform and become phagocytic, as we observed a tight correlation between astrocyte transformation and Beat-Va_M_ neurite fragmentation. We note that the fragmentation we observed when EcR signaling was blocked in Beat-Va_M_ neurons occurred primarily in smaller diameter Beat-Va_M_ neurites. Perhaps the architectural integrity and microtubule and actin content, of large versus small diameter of a neuronal process makes fine neurite processes more susceptible to remodeling.

Beyond physical forces, what astrocyte-derived signals might promote Beat-Va_M_ neuron local pruning? Astrocytes may release molecular cues onto neurites as they transform into phagocytes that initiate neurite remodeling. We explored the role of the previously identified glial-secreted TGFβ ligand Myoglianin in Beat-Va_M_ neuron local pruning. In other cell types, Myoglianin signaling from glia is known to activate the expression of EcR in neurons, thereby establishing their competence to prune in response to ecdysone (Awasaki, Huang et al. 2011, Yu, Gutman et al. 2013, Hakim, Yaniv et al. 2014, Wang, Lee et al. 2019). Multiple lines of evidence argue against such a role for Myoglianin in Beat-Va_M_ neuron local pruning. First, we found that while blockade of pan-glial release of Myoglianin can suppress Beat-Va_M_ neuron local pruning, depletion of Myoglianin from astrocytes did not block Beat-Va_M_ neuron local pruning. Second, we found that Myoglianin depletion led to decreased EcR expression in astrocytes and a failure of astrocytes to transform, arguing that Myoglianin from non-astrocytic glia drives astrocyte expression of EcR and, in turn, transformation into phagocytes. Finally, we find that blockade of EcR signaling autonomously in Beat-Va_M_ neurons has only minimal effects on Beat-Va_M_ local pruning of neurites. We, therefore, speculate that astrocytes activate Beat-Va_M_ neuron local pruning either through secretion of a different signaling factor, through the physical force astrocytes generate during transformation into phagocytes, or both.

We further demonstrated that Beat-Va_L_ neurons die in a caspase and steroid hormone-dependent fashion, similar to other neuronal cell types that undergo cell death at metamorphosis (Choi 2006, Lee, Sehgal et al. 2013). However, we found that the Hox gene *Abd-B* controls the segment-specific patterns of Beat-Va_L_ neuron cell death, with the three Abd-B+ posterior Beat-Va_L_ cells undergoing cell death, while the anterior two Abd-B-negative cells survive. Knocking down *Abd-B* in Beat-Va neurons blocked caspase activation and cell death in the posterior cells, while overexpressing *Abd-B* in Beat-Va neurons drove cell death in the two normally surviving anterior Beat-Va_L_ cells. Notably, neither overexpression nor knockdown of *Abd-B* had any noticeable effect on Beat-Va_M_ neuron local pruning despite their having a similar segmentally restricted pattern of Abd-B expression in segments A5-7. We interpret these data to mean that Abd-B confers positional identity that leads to cell death in the appropriate Beat-Va_L_ cells.

Hox genes such as *Abd-B* can function as a pro-apoptotic or anti-apoptotic molecules. In some cases, Abd-B can drive caspase-mediated cell death through direct regulation of gene expression. Abd-B-driven chromatin remodeling can lead to exposure of activator genes *grim, reaper,* and *hid*, thereby enabling activation of apoptotic death (Arya, Sarkissian et al. 2015). Abd-B can also physically bind the transcription co-factor Dachshund, and these factors have been shown to work cooperatively to induce cell death in some embryonic neurons (Clarembaux-Badell, Baladrón-de-Juan et al. 2022). In other cases, for instance, in Beat-Va_M_ neurons, Abd-B expression clearly does not regulate cell death at all, presumably because pro-or anti-apoptotic functions of Abd-B are regulated by cell type-specific co-factors. Finally, additional studies of the Abd-B ortholog, HOXA10A, have revealed that hox genes can modify cell death through production of long non-coding RNAs and other post transcriptional modifications during cancer metastasis (Chen, Kan et al. 2022). Whether and how these mechanisms might relate to the role of Abd-B in driving segment-specific cell death is an interesting question for the future.

## Methods

### Resource Table

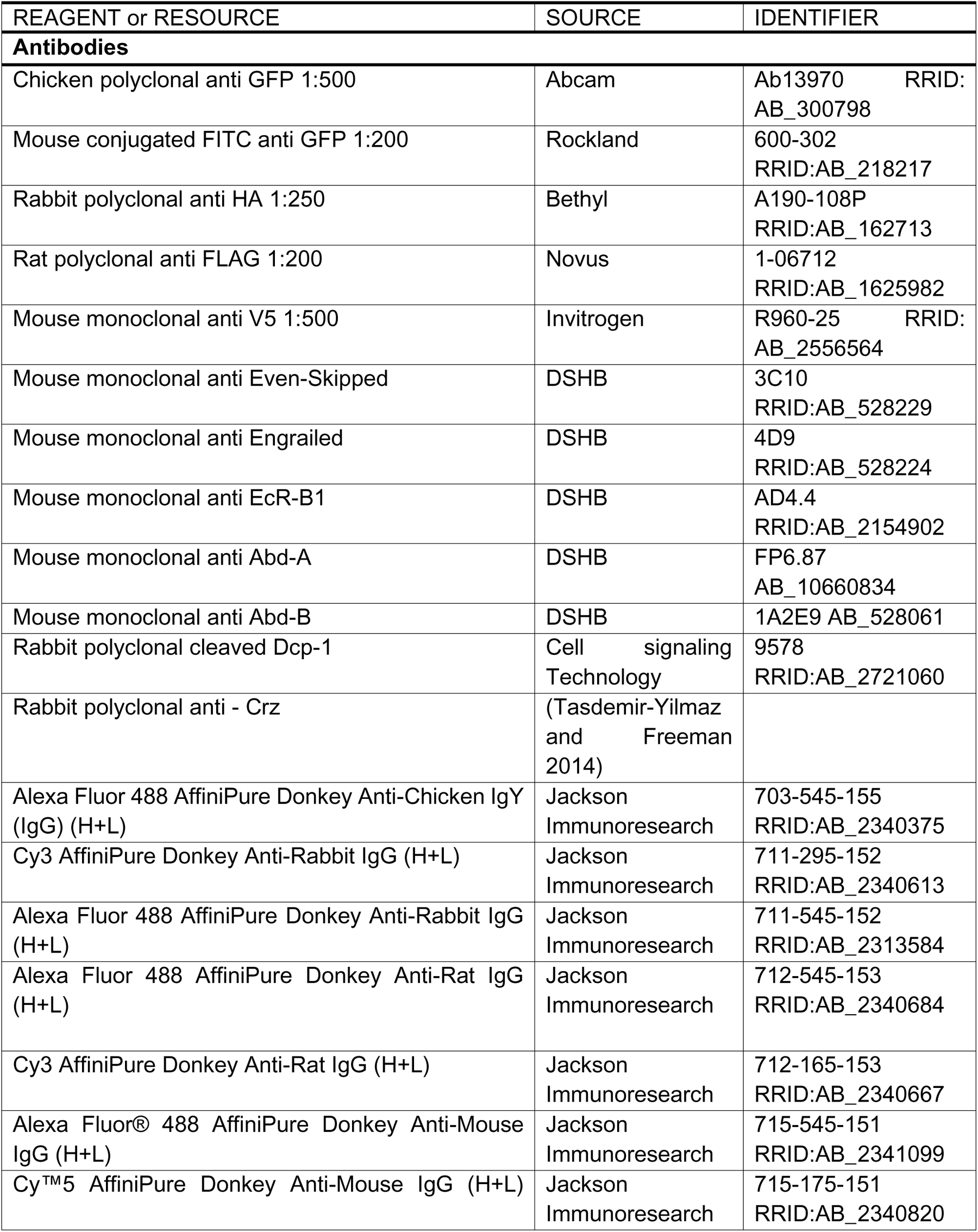
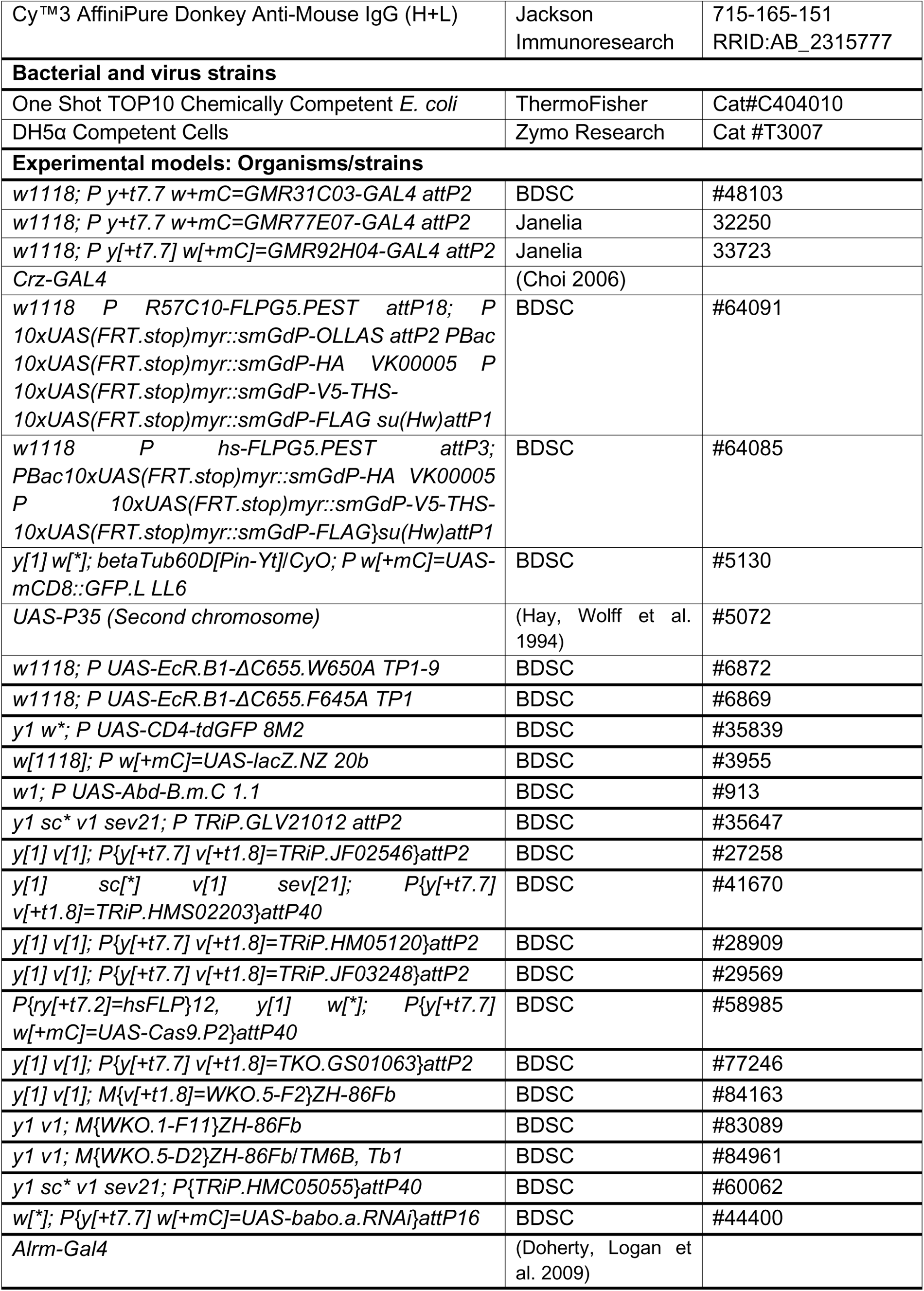
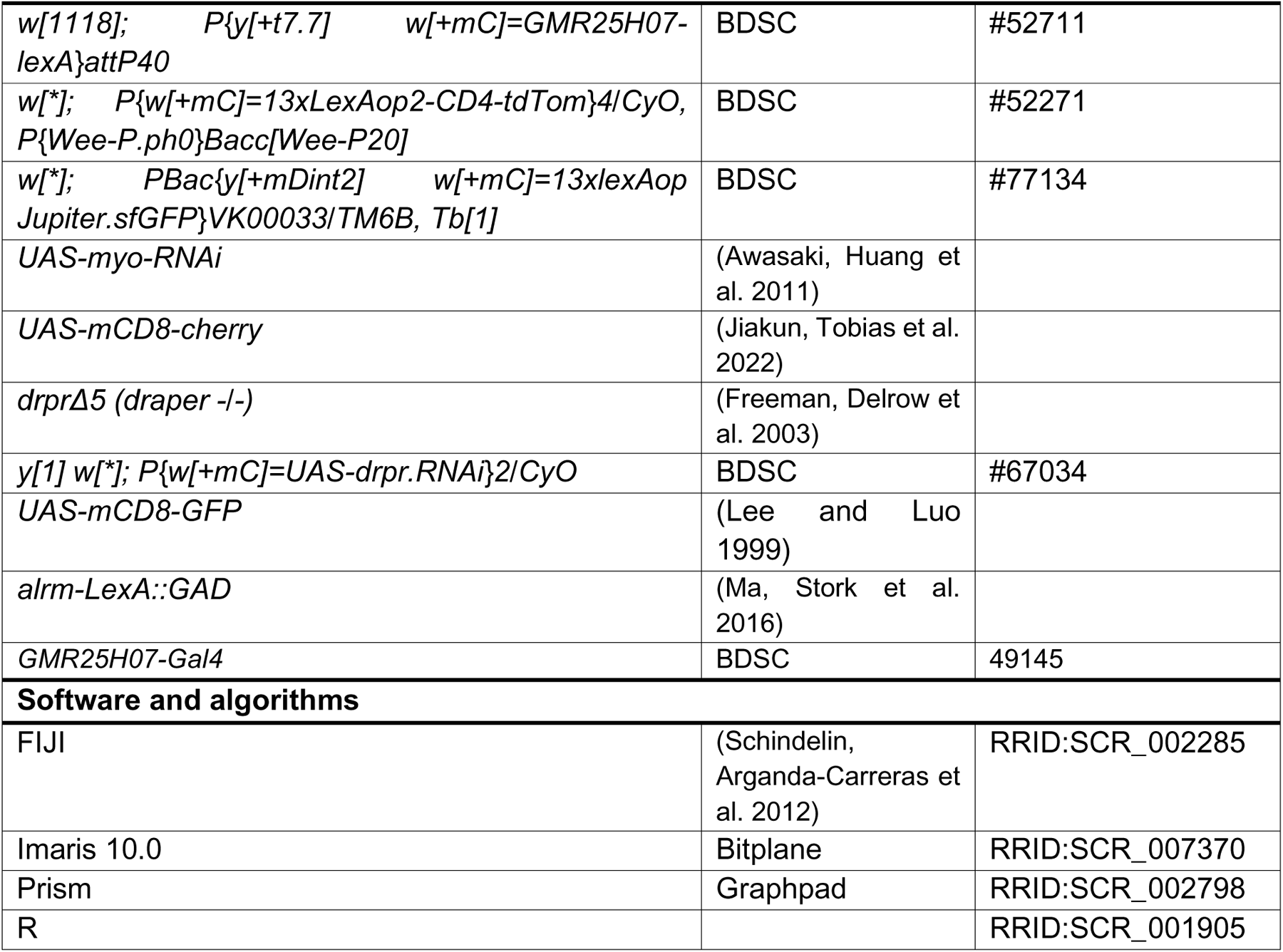

### Drosophila Genetics

*Drosophila melanogaster* were reared using standard laboratory conditions. All experiments were conducted at 25°C unless otherwise explicitly noted. A complete list of all fly strains used in this study can be found in the key resource table.

### Immunohistochemistry

Dissection and immunostaining of larval and pupal fly brains were performed according to an adapted FlyLight protocol (Jenett, Gerald et al. 2012). Larval brains were dissected in cold PBST (phosphate buffered saline, Invitrogen) with 0.1% Triton X-100 and fixed at 4% paraformaldehyde (PFA, Electron Microscopy Sciences) for 20 minutes at room temperature. Fixed brains were washed three times with PBS while nutating at room temperature for 10-15 minutes per wash. Primary and secondary antibodies were diluted in PBST (0.1% Triton X-100) and incubated with samples at 4° C for 24-48 hours. Washes after antibody incubations were performed at room temperature with PBS for 3 X 15 minutes. Samples were mounted in VECTASHIELD antifade mounting medium (Vector Laboratories) and stored at 4° C until imaging.

### Image Analysis and Processing

Imaging was performed using a Zeiss LSM 880 with Airyscan. Confocal z-stacks were acquired using the optimal z-interval and 0.05-0.09 µm/pixel resolution with a 40x/1.3 Plan-Apochromat oil objective. Images were Airyscan processed, image tiles stitched with a 6-8% overlap when necessary and converted into IMARIS format for 3D analysis. 3D rendering was performed with IMARIS; single image or z-stack projections were analyzed in ImageJ.

### Image Analysis

To quantify individual neuronal morphologies, we used the Surface module in IMARIS 10 (Bitplane) with the Filament module. The module was trained on example images through iterative machine learning and then the algorithm was applied to all images. The first 100 µm of each neuron after it crossed the midline was analyzed for both total filament length and number of branch points.

### Data Analysis and Statistics

All statistical analyses were carried out using GraphPad Prism 9; statistical details, p values, and sample sizes are indicated in the figure legends. Any comparison between two parametric data sets that only had one independent variable was conducted with a student t-test. Any comparison between more than two parametric data sets that only had one independent variable was conducted with a One-way ANOVA with the Sidak test for multiple comparisons. Any comparison between more than two non-parametric data sets that only had one independent variable was conducted with a One-way ANOVA with the Kruskal-Wallis test for multiple comparisons. Any comparison between more than two parametric data sets that had two independent variables was conducted with a Two-way ANOVA with the Sidak test for multiple comparisons.

### Generating *BeatVa-LexA* Stock

*BeatVa-LexA* lines were made with Gateway cloning. The 3471 base pair enhancer region was cloned with primer sequences provided on the Janelia Flylight page. They were cloned into the *pBPnlsLexAGADflUw* vector (#26232 Addgene) and verified through Sanger sequencing (Genewiz). Injections were site directed using the *y^1^ w^67c23^; P[CaryP attP2]* line (BL8622), *y^1^ w^67c23^ P[CaryP attP18]* (BL 32107) line and *y^1^ w^67c23^; P[CaryP attP40]* line. The attp18 insertion was used in all above experiments. The line was verified by comparing GFP expression generation from the *Beat-LexA* line driving a *LexAop-GFP* to a *BeatVa-Gal4* line driving a *UAS-mCD8::cherry*.

### Generating *LexAop-EcR^DN^* Stock

The *LexAop-EcR^DN^* construct was created by amplifying DNA from flies containing *UAS-EcR.B1DC655.W650A* (Bloomington Drosophila Stock Center). The amplification was carried out using the EcR^DN^ Kozak *AgeI* F primer (cccccaACCGGTcaaaacATGAAGCGGCGCTGGTCGAACA) and EcR^DN-W650A^ *XbaI* R primer (cccaatctagaCTAGATGGCATGAACGTCGGCGA). The resulting PCR product was digested with AgeI and XbaI (NEB) and then ligated into the *pattB-13xLexAop2EGFP* plasmid (Coutinho-Budd, Sheehan et al. 2017), with GFP removed via *AgeI* and *XbaI* digestion. The final construct, *pattB-13xLexAop2-EcR^W650A^*, was generated through ligation using T4 ligase (NEB), and the transformation was carried out in *DH5α* cells (Zymo). The construct’s sequence was verified through Sanger sequencing (Genewiz), and it was utilized for attp154 landing site-specific integration on the 2nd chromosome (Bestgene).

---

**Supplemental Figure 1:**
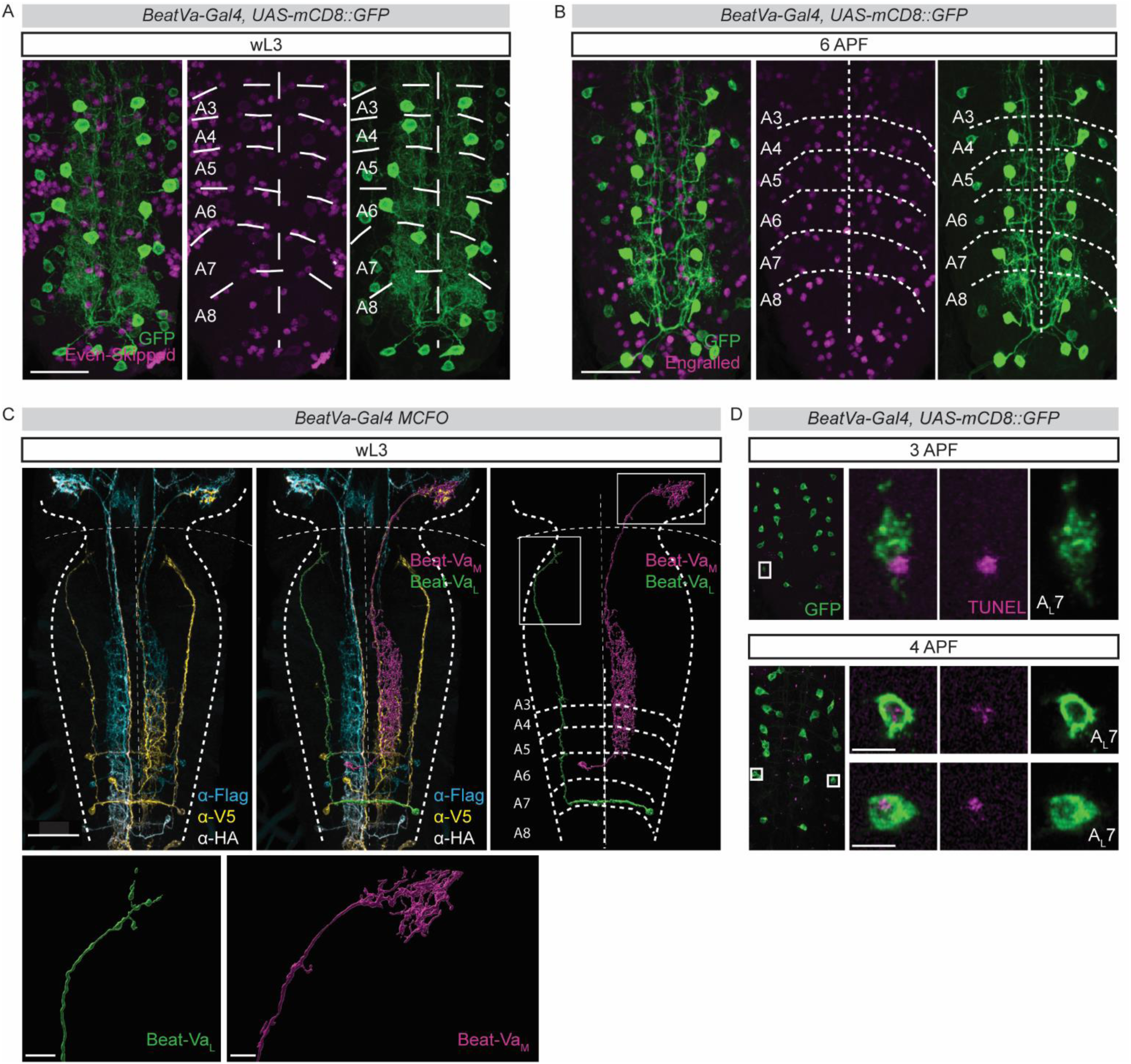
Segmental characterization of Beat-Va_M_ and Beat-Va_L_ neurons. **A)** VNC at wL3 with genetic mCD8::GFP labeling of Beat-Va neurons (green) and anti-Even-Skipped staining (magenta) to label the segments of the ventral nerve cord. Segments A3-A8 are denoted by tracing the Even-Skipped staining and then superimposed onto Beat-Va neurons to define segmental positions. Scale bar is 20 microns. **B)** VNC at 6APF with genetic GFP labeling of Beat-Va neurons (green) and anti-Engrailed staining (magenta) to label the segments of the ventral nerve cord, showing the segmental positions persist into metamorphosis. Scale bars are 20 microns. **C)** Using segmental information and MCFO we can render single cells in Imaris and overlay positional information. Boxed areas are Beat-Va_L_ and Beat-Va_M_ termini and are displayed in high magnification (Scale bars are 10 microns). CNS is outlined and boundary between VNC and brain lobes is marked with a dashed line. **D**) Z-projection of ventral nerve cords with Beat-Va neurons labeled genetically with mCD8::GFP (green) and then subjected to TUNEL staining (magenta) to detect cells undergoing apoptotic cell death at 3APF and 4APF. Boxed areas containing TUNEL positive lateral cells are magnified and segmental position is noted. Scale bar is 5 microns.

**Supplemental Figure 2:**
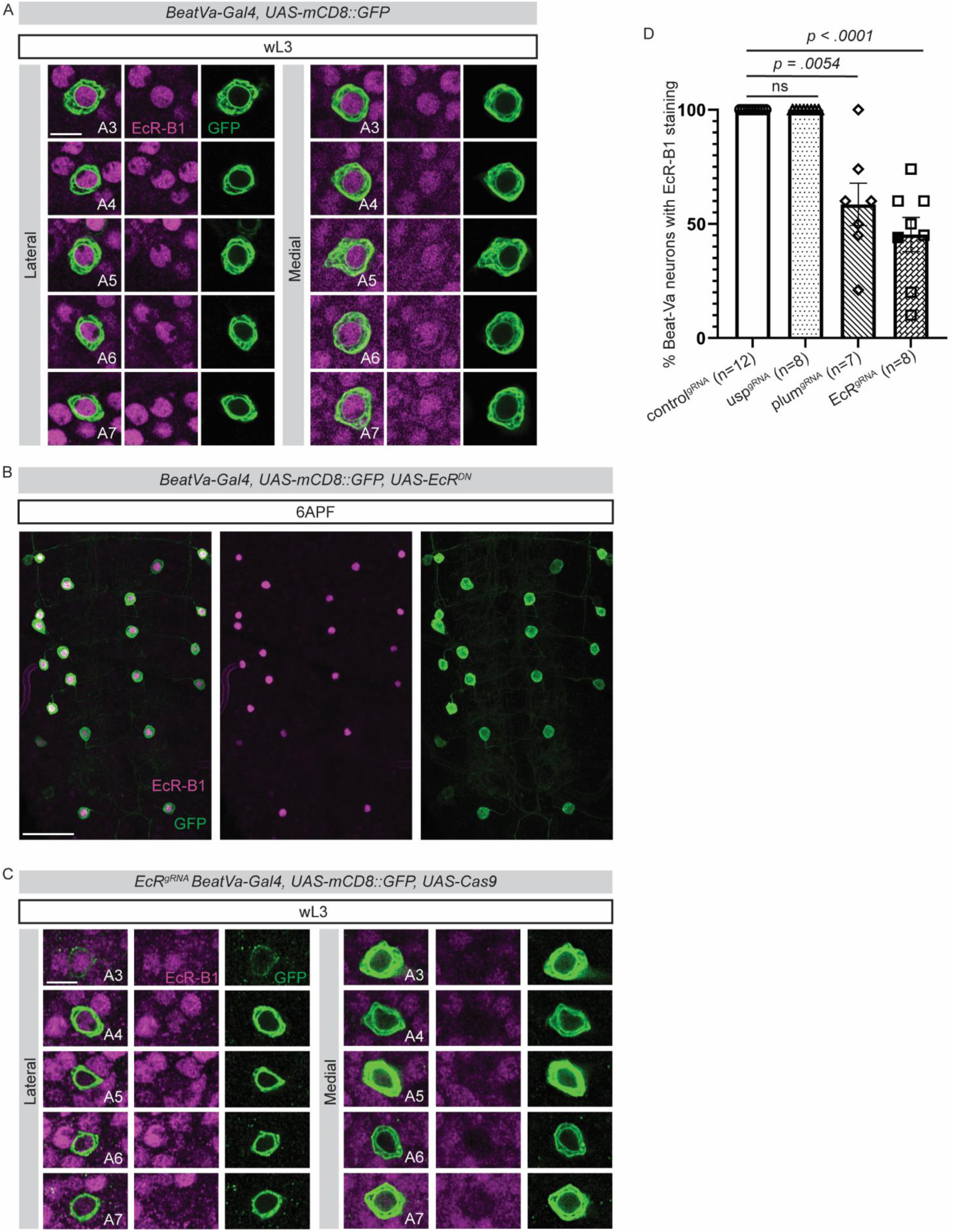
Manipulation of EcR-B1 expression in Beat-Va neurons. **A)** Beat-Va neurons at wL3 labeled CD8::GFP (green) stained with the EcR B1 Receptor (magenta) showing the colocalization of all lateral and medial cell bodies with the receptor. **B)** Beat-Va neurons at 6APF labeled CD8::GFP (green) and expressing a UAS drive EcR^DN^. Stained with the EcR B1 Receptor (magenta) to identify cells that are expressing the *UAS-EcR^DN^.* There are typically not EcR-B1 positive cells at this time. **C)** Beat-Va neurons at wL3 labeled CD8::GFP (green) and expressing *UAS-Cas9* in a gRNA EcR animal. Stained with the EcR-B1 Receptor (magenta) to evaluate if cells lose EcR-B1 expression. **D)** Quantification of gRNA against *usp*, *plum, EcR* on Beat-Va EcR-B1 expression when compared with control. One-Way ANOVA, Kruskal-Wallis test for multiple comparisons.

**Supplemental Figure 3:**
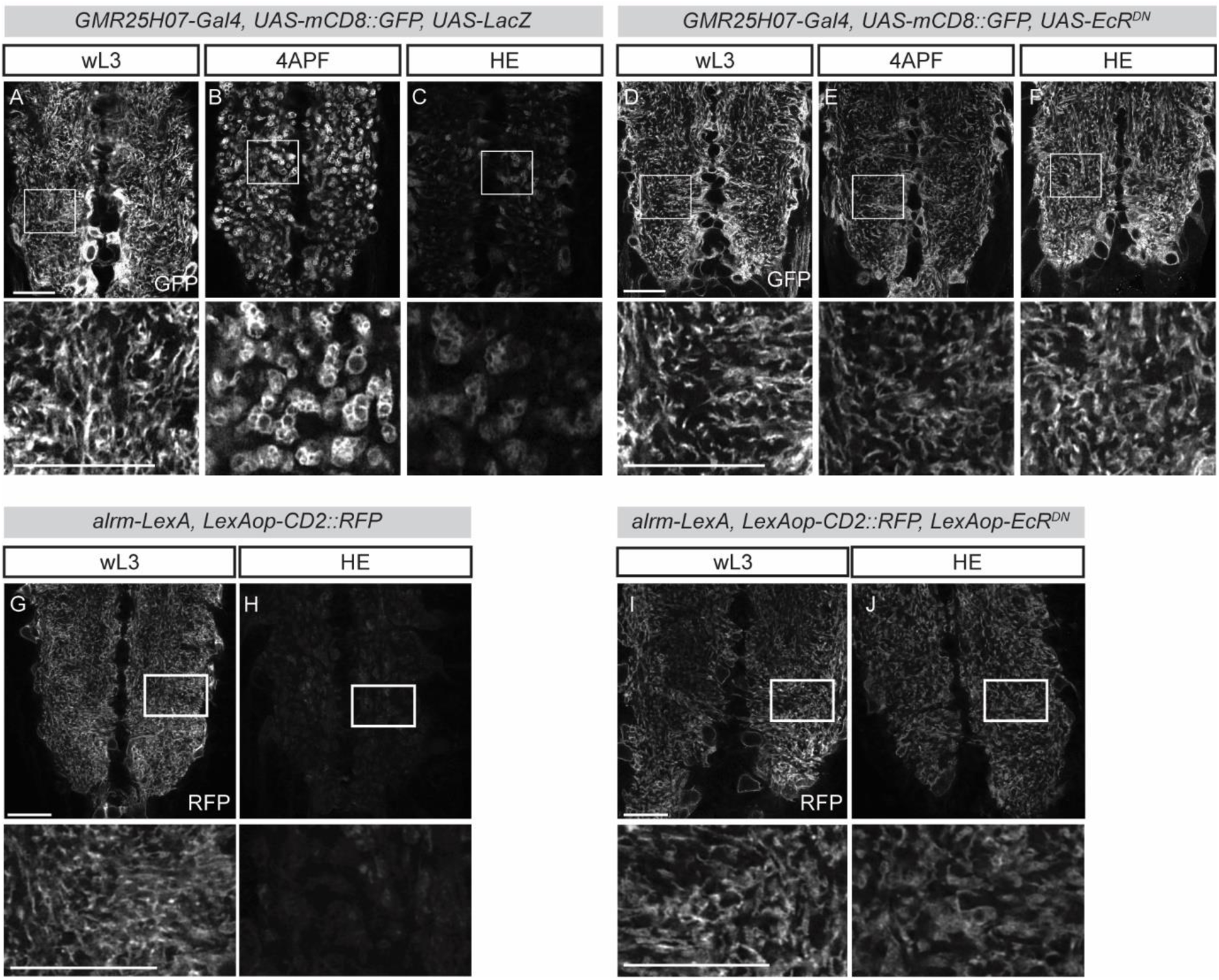
Cell specific expression of EcR^DN^ blocks astrocyte transformation. **A)** Astrocytes labeled with *GMR25H07-Gal4* (a strong astrocyte Gal4 line) crossed to CD8::GFP at wL3 showing their “wispy” morphology, **B)** 4APF showing the phagolysomes that astrocytes display (Kang 2023), and **C)** HE where astrocytes are normally only faintly detectable. **D)** Astrocytes expressing a *UAS-EcR^DN^*, labeled with mCD8::GFP at wL3 showing their “wispy” morphology, which is retained at **E)** 4APF, and **F)** HE, indicating a failure to transform. **G)** *arm-LexA, LexAop-mCD8::RFP* at wL3 where normal astrocyte morphology is apparent, and **H)** HE when astrocytes are no longer visible. **I)** When *alrm-LexA* drives a *LexAop-EcR^DN^*in addition to *LexAop-mCD8::RFP,* the astrocytes appear normal at wL3, **J)** but then fail to transform at HE. Scale bars are 20 microns.

**Supplemental Figure 4:**
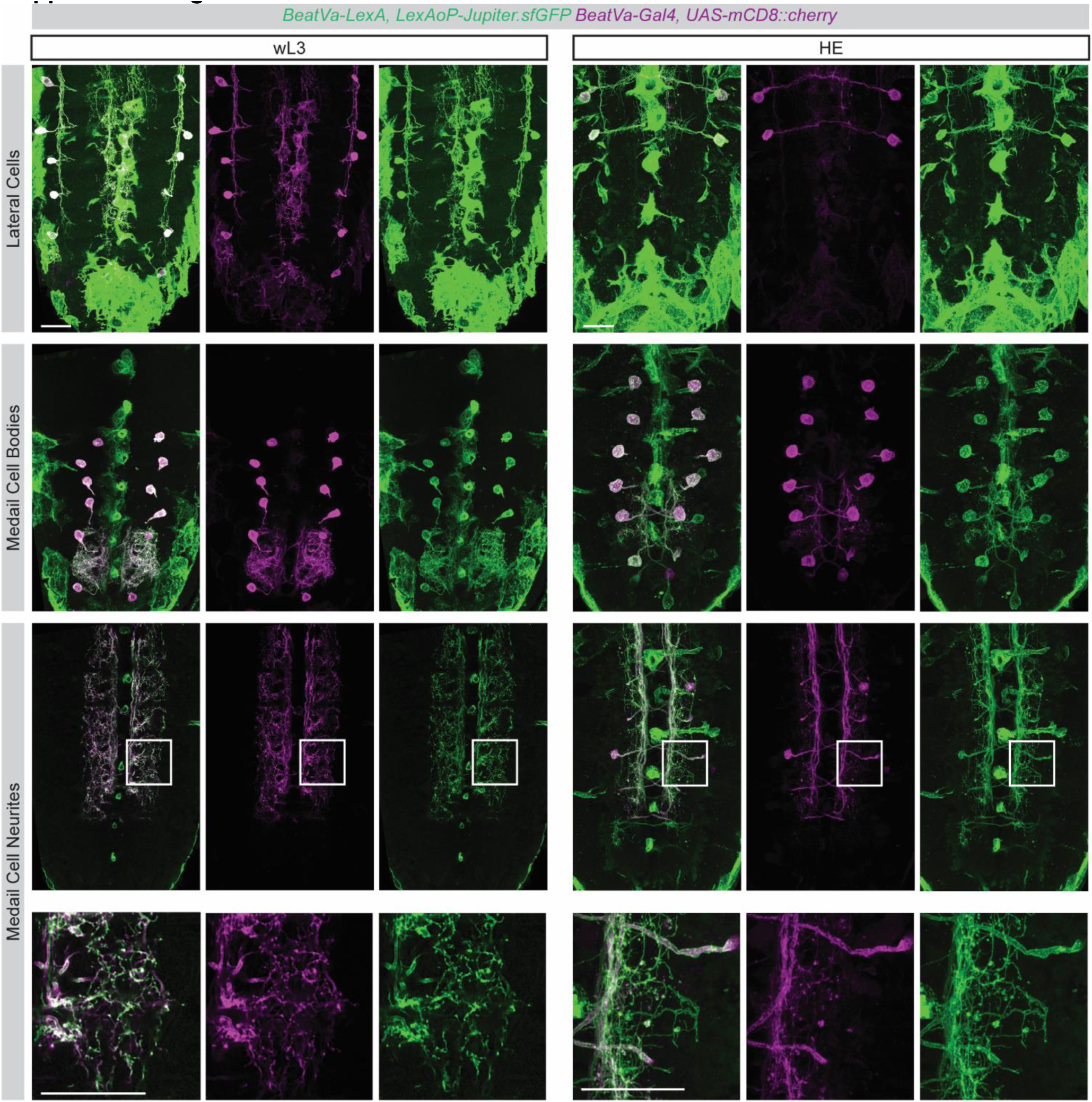
Verification of newly constructed *BeatVa-LexA* line. Animals carrying the *BeatVa-LexA* construct were crossed to animals carrying a *LexAop-Jupiter.sfGFP* (a GFP construct that localized to microtubule networks and had been reported to label neurite processes well (Karpova, Bobinnec et al. 2006)) to genetically label any cells where the *BeatVa-LexA* line was expressed. These animals were then crossed to an existing stock that carried both the original *BeatVa-Gal4* construct and a *UAS-mCD8::cherry*. Good signal overlap between the GFP and Cherry fluorophores indicates the *BeatVa-LexA* and *BeatVa-Gal4* lines labeled the same population of neurons in addition to labeling a second population of non-neuronal cells (likely surface glia). Scale bars are 20 microns.

**Supplemental Figure 5:**
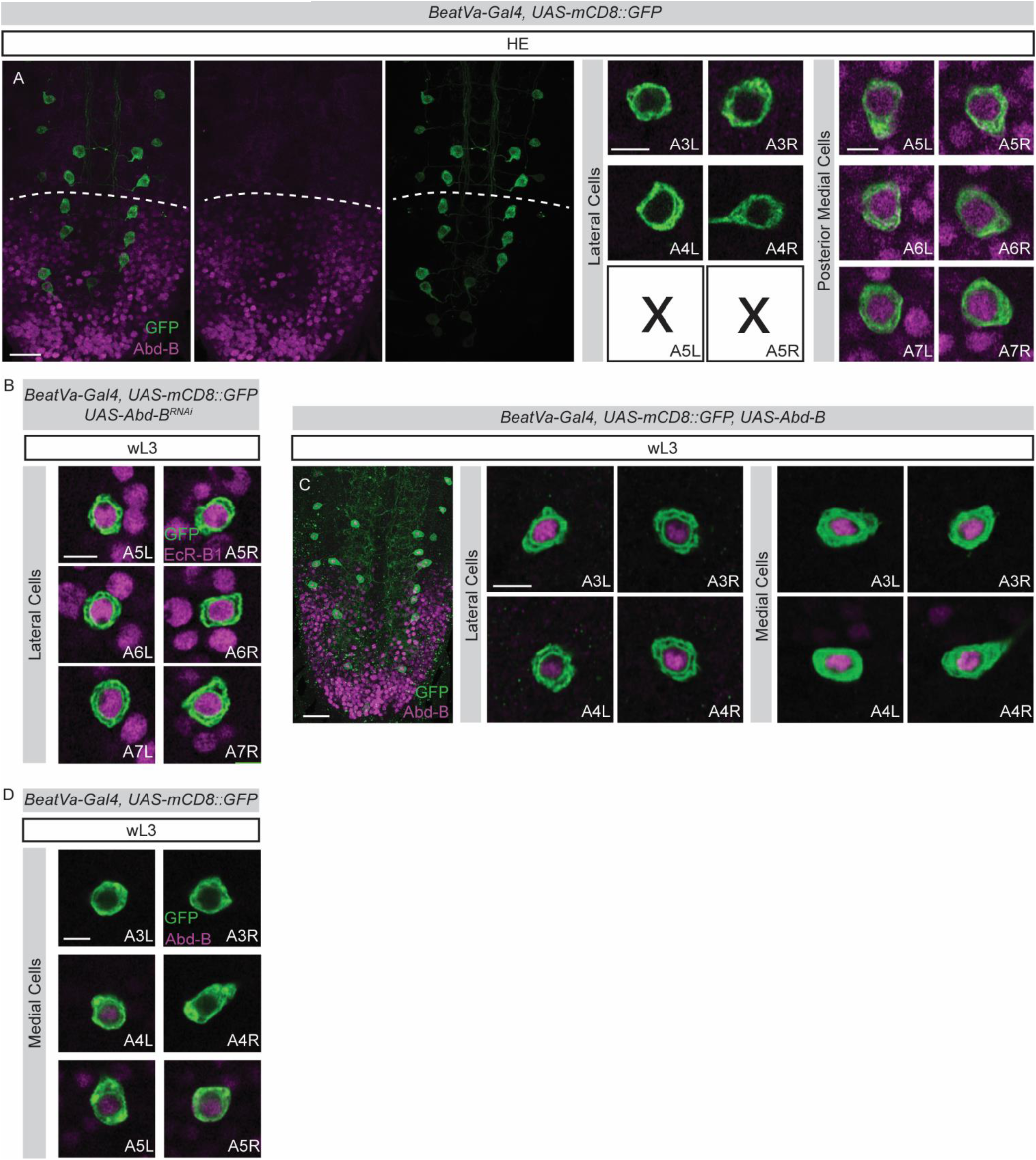
Abd-B expression in Beat-Va neurons. **A)** Beat-Va neurons at HE labeled with mCD8::GFP stained with anti-Abd-B. Abd-B expression persists to HE. The lateral A5 (Abd-B +) cells undergo cell death while the A3-4 cells survive and do not express Abd-B at HE. The A5-7 medial cells continue to express Abd-B at HE. Left and right hemisegment denoted by “L” and “R”. Dashed white line superimposed on Abd-B boundary. **B)** Beat-Va neurons labeled with mCD8::GFP and driving *UAS-Abd-B^RNAi^*at wL3 stained with the EcR-B1 showing the continued colocalization of EcR-B1 in the absence of Abd-B. **C)** Beat-Va neurons at wL3 labeled with mCD8::GFP and crossed to *UAS-Abd-B* and stained with anti-Abd-B. The two most anterior lateral and medial cell bodies are magnified and show Abdominal-B expression, which is not usually present in these cells, confirming overexpression was successful. **D)** Three most anterior medial Beat-Va neurons at wL3 labeled with CD8::GFP stained with Abd-B. Shows Abd-B is present in some medial cells even though they do not undergo cell death. Scale bar is 5 microns in all images with a single cell body and 20 microns in all other images.

## Acknowledgements

We would like to acknowledge all members of the Freeman lab for thoughtful discussions and suggestions throughout this project. We acknowledge the fly community for generous sharing of reagents. The authors declare no competing financial interests.

This study was supported by National Science Foundation GFRP Fellowship DGE-1448072 (KSL), Damon Runyon Fellowship DRG 2329-18 (YK), and National Institutes of Health grants K99 NS126642-01 (YK) 3R01NS059991-14S1 (MRF)

## Author Contributions

Conceptualization, K.S. Lehmann, Y. Kang and M.R. Freeman; Data Curation, K.S. Lehmann, M. Hupp, A. Jefferson, Y.C. Chang, A. Sheehan; Formal Analysis, K.S. Lehmann, Y. Kang, A. Sheehan; Funding Acquisition, K.S. Lehmann, Y. Kang, M.R. Freeman; Investigation, K.S. Lehmann, M. Hupp, A. Jefferson, A. Sheehan Y. Kang; Methodology, K.S. Lehmann, Y. Kang, M.R. Freeman; Resources; M.R. Freeman; Supervision, Y. Kang, M.R. Freeman; Validation, M. Hupp, A. Jefferson; Visualization, K.S. Lehmann; Writing—Original Draft, K.S. Lehmann and M.R. Freeman; Writing—Reviewing & Editing, K.S. Lehmann, A. Jefferson, Y. Kang, M.R. Freeman.

## References

Stevens, B., N. J. Allen, L. E. Vazquez, G. R. Howell, K. S. Christopherson, N. Nouri, K. D. Micheva, A. K. Mehalow, A. D. Huberman, B. Stafford, A. Sher, Alan, J. D. Lambris, S. J. Smith, S. W. M. John and B. A. Barres (2007). “The Classical Complement Cascade Mediates CNS Synapse Elimination.” Cell 131(6): 1164–1178.

Williams, D. W. and J. W. Truman (2005). “Cellular mechanisms of dendrite pruning in Drosophila: insights from in vivo time-lapse of remodeling dendritic arborizing sensory neurons.” Development 132(16): 3631–3642.

Stanfield, B. B., D. D. M. O’Leary and C. Fricks (1982). “Selective collateral elimination in early postnatal development restricts cortical distribution of rat pyramidal tract neurones.” Nature 298(5872): 371–373.

Karcavich, R. and C. Q. Doe (2005). “Drosophila neuroblast 7-3 cell lineage: A model system for studying programmed cell death, Notch/Numb signaling, and sequential specification of ganglion mother cell identity.” Journal of Comparative Neurology 481(3): 240–251.

Atz, M. E., B. Rollins and M. P. Vawter (2007). “NCAM1 association study of bipolar disorder and schizophrenia: polymorphisms and alternatively spliced isoforms lead to similarities and differences.” Psychiatr Genet 17(2): 55–67.

Feinberg, I. (1982). “Schizophrenia: caused by a fault in programmed synaptic elimination during adolescence?” J Psychiatr Res 17(4): 319–334.

Ishizuka, K., Y. Fujita, T. Kawabata, H. Kimura, Y. Iwayama, T. Inada, Y. Okahisa, J. Egawa, M. Usami, I. Kushima, Y. Uno, T. Okada, M. Ikeda, B. Aleksic, D. Mori, T. Someya, T. Yoshikawa, N. Iwata, H. Nakamura, T. Yamashita and N. Ozaki (2017). “Rare genetic variants in CX3CR1 and their contribution to the increased risk of schizophrenia and autism spectrum disorders.” Transl Psychiatry 7(8): e1184.

Neniskyte, U. and C. T. Gross (2017). “Errant gardeners: glial-cell-dependent synaptic pruning and neurodevelopmental disorders.” Nature Reviews Neuroscience 18(11): 658–670.

Sekar, A., A. R. Bialas, H. de Rivera, A. Davis, T. R. Hammond, N. Kamitaki, K. Tooley, J. Presumey, M. Baum, V. Van Doren, G. Genovese, S. A. Rose, R. E. Handsaker, M. J. Daly, M. C. Carroll, B. Stevens and S. A. McCarroll (2016). “Schizophrenia risk from complex variation of complement component 4.” Nature 530(7589): 177–183.

Selemon, L. D., G. Rajkowska and P. S. Goldman-Rakic (1995). “Abnormally high neuronal density in the schizophrenic cortex. A morphometric analysis of prefrontal area 9 and occipital area 17.” Arch Gen Psychiatry 52(10): 805–818; discussion 819-820.

Selemon, L. D., G. Rajkowska and P. S. Goldman-Rakic (1998). “Elevated neuronal density in prefrontal area 46 in brains from schizophrenic patients: application of a three-dimensional, stereologic counting method.” J Comp Neurol 392(3): 402–412.

Winchester, C. L., H. Ohzeki, D. A. Vouyiouklis, R. Thompson, J. M. Penninger, K. Yamagami, J. D. Norrie, R. Hunter, J. A. Pratt and B. J. Morris (2012). “Converging evidence that sequence variations in the novel candidate gene MAP2K7 (MKK7) are functionally associated with schizophrenia.” Human molecular genetics 21(22): 4910–4921.

Dorothy, Emily, Amanda, R. Koyama, Alan, R. Yamasaki, Richard, Michael, Ben and B. Stevens (2012). “Microglia Sculpt Postnatal Neural Circuits in an Activity and Complement-Dependent Manner.” Neuron 74(4): 691–705.

Luo, L. and D. D. O’Leary (2005). “Axon retraction and degeneration in development and disease.” Annu Rev Neurosci 28: 127–156.

Neukomm, L. J. and M. R. Freeman (2014). “Diverse cellular and molecular modes of axon degeneration.” Trends Cell Biol 24(9): 515–523.

Riccomagno, M. M. and A. L. Kolodkin (2015). “Sculpting neural circuits by axon and dendrite pruning.” Annu Rev Cell Dev Biol 31: 779–805.

Hutchins, J. B. and S. W. Barger (1998). “Why neurons die: cell death in the nervous system.” Anat Rec 253(3): 79–90.

Oppenheim, R. W. (1985). “Naturally occurring cell death during neural development.” Trends in Neurosciences 8: 487–493.

Southwell, D. G., M. F. Paredes, R. P. Galvao, D. L. Jones, R. C. Froemke, J. Y. Sebe, C. Alfaro-Cervello, Y. Tang, J. M. Garcia-Verdugo, J. L. Rubenstein, S. C. Baraban and A. Alvarez-Buylla (2012). “Intrinsically determined cell death of developing cortical interneurons.” Nature 491(7422): 109–113.

Petersen, C. C. H. (2019). “Sensorimotor processing in the rodent barrel cortex.” Nature Reviews Neuroscience 20(9): 533–546.

Gunner, G., L. Cheadle, K. M. Johnson, P. Ayata, A. Badimon, E. Mondo, M. A. Nagy, L. Liu, S. M. Bemiller, K. W. Kim, S. A. Lira, B. T. Lamb, A. R. Tapper, R. M. Ransohoff, M. E. Greenberg, A. Schaefer and D. P. Schafer (2019). “Sensory lesioning induces microglial synapse elimination via ADAM10 and fractalkine signaling.” Nat Neurosci 22(7): 1075–1088.

Hoeppner, D. J., M. O. Hengartner and R. Schnabel (2001). “Engulfment genes cooperate with ced-3 to promote cell death in Caenorhabditis elegans.” Nature 412(6843): 202–206.

Reddien, P. W., S. Cameron and H. R. Horvitz (2001). “Phagocytosis promotes programmed cell death in C. elegans.” Nature 412(6843): 198–202.

Marín-Teva, J. L., I. Dusart, C. Colin, A. Gervais, N. van Rooijen and M. Mallat (2004). “Microglia promote the death of developing Purkinje cells.” Neuron 41(4): 535–547.

Lee, T., A. Lee and L. Luo (1999). “Development of the Drosophila mushroom bodies: sequential generation of three distinct types of neurons from a neuroblast.” Development 126(18): 4065–4076.

Lee, T., S. Marticke, C. Sung, S. Robinow and L. Luo (2000). “Cell-autonomous requirement of the USP/EcR-B ecdysone receptor for mushroom body neuronal remodeling in Drosophila.” Neuron 28(3): 807–818.

Choi, Y. J. (2006). “Programmed cell death mechanisms of identifiable peptidergic neurons in Drosophila melanogaster.” 133(11): 2223–2232.

Watts, R. J., E. D. Hoopfer and L. Luo (2003). “Axon Pruning during Drosophila Metamorphosis: Evidence for Local Degeneration and Requirement of the Ubiquitin-Proteasome System.” Neuron 38(6): 871–885.

Kirilly, D., Y. Gu, Y. Huang, Z. Wu, A. Bashirullah, B. C. Low, A. L. Kolodkin, H. Wang and F. Yu (2009). “A genetic pathway composed of Sox14 and Mical governs severing of dendrites during pruning.” Nature neuroscience 12(12): 1497–1505.

Zhang, H., Y. Wang, Jack Jing L. Wong, K.-L. Lim, Y.-C. Liou, H. Wang and F. Yu (2014). “Endocytic Pathways Downregulate the L1-type Cell Adhesion Molecule Neuroglian to Promote Dendrite Pruning in Drosophila.” Developmental Cell 30(4): 463–478.

Truman, J. W., W. S. Talbot, S. E. Fahrbach and D. S. Hogness (1994). “Ecdysone receptor expression in the CNS correlates with stage-specific responses to ecdysteroids during Drosophila and Manduca development.” Development (Cambridge, England) 120(1): 219–234.

Schubiger, M., A. A. Wade, G. E. Carney, J. W. Truman and M. Bender (1998). “Drosophila EcR-B ecdysone receptor isoforms are required for larval molting and for neuron remodeling during metamorphosis.” Development 125(11): 2053–2062.

Schubiger, M., S. Tomita, C. Sung, S. Robinow and J. W. Truman (2003). “Isoform specific control of gene activity in vivo by the Drosophila ecdysone receptor.” Mechanisms of Development 120(8): 909–918.

Thummel, C. S. (1996). “Flies on steroids — Drosophila metamorphosis and the mechanisms of steroid hormone action.” Trends in Genetics 12(8): 306–310.

Talbot, W. S., E. A. Swyryd and D. S. Hogness (1993). “Drosophila tissues with different metamorphic responses to ecdysone express different ecdysone receptor isoforms.” Cell 73(7): 1323–1337.

Riddiford, L. M. and J. W. Truman (1993). “Hormone receptors and the regulation of insect metamorphosis.” American Zoologist 33(3): 340–347.

Koelle, M. R., W. S. Talbot, W. A. Segraves, M. T. Bender, P. Cherbas and D. S. Hogness (1991). “The drosophila EcR gene encodes an ecdysone receptor, a new member of the steroid receptor superfamily.” Cell 67(1): 59–77.

Pinto-Teixeira, F., N. Konstantinides and C. Desplan “Programmed cell death acts at different stages of Drosophila neurodevelopment to shape the central nervous system.” (1873–3468 (Electronic)).

Yamaguchi, Y. and M. Miura (2015). “Programmed Cell Death in Neurodevelopment.” Developmental Cell 32(4): 478–490.

Marchetti, G. and G. Tavosanis (2017). “Steroid Hormone Ecdysone Signaling Specifies Mushroom Body Neuron Sequential Fate via Chinmo.” Current Biology 27(19): 3017–3024.e3014.

Hara, Y., K. Hirai, Y. Togane, H. Akagawa, K. Iwabuchi and H. Tsujimura (2013). “Ecdysone-dependent and ecdysone-independent programmed cell death in the developing optic lobe of Drosophila.” Developmental biology 374(1): 127–141.

Winbush, A. and J. C. Weeks (2011). “Steroid-triggered, cell-autonomous death of a Drosophila motoneuron during metamorphosis.” Neural Dev 6: 15.

Kuo, C. T., L. Y. Jan and Y. N. Jan (2005). “Dendrite-specific remodeling of Drosophila sensory neurons requires matrix metalloproteases, ubiquitin-proteasome, and ecdysone signaling.” Proceedings of the National Academy of Sciences of the United States of America 102(42): 15230–15235.

Tasdemir-Yilmaz, O. E. and M. R. Freeman (2014). “Astrocytes engage unique molecular programs to engulf pruned neuronal debris from distinct subsets of neurons.” Genes Dev 28(1): 20–33.

Lee, G., R. Sehgal, Z. Wang, S. Nair, K. Kikuno, C.-H. Chen, B. Hay and J. H. Park (2013). “Essential role of grim-led programmed cell death for the establishment of corazonin-producing peptidergic nervous system during embryogenesis and metamorphosis in Drosophila melanogaster.” Biology Open 2(3): 283–294.

Lee, G., R. Sehgal, Z. Wang and J. H. Park (2019). “Ultraspiracle-independent anti-apoptotic function of ecdysone receptors is required for the survival of larval peptidergic neurons via suppression of grim expression in Drosophila melanogaster.” Apoptosis : an international journal on programmed cell death 24(3-4): 256–268.

Lee, G., Z. Wang, R. Sehgal, C.-H. Chen, K. Kikuno, B. Hay and J. H. Park (2011). “Drosophila caspases involved in developmentally regulated programmed cell death of peptidergic neurons during early metamorphosis.” Journal of Comparative Neurology 519(1): 34–48.

Wang, Z., G. Lee, R. Vuong and J. H. Park (2019). “Two-factor specification of apoptosis: TGF-β signaling acts cooperatively with ecdysone signaling to induce cell- and stage-specific apoptosis of larval neurons during metamorphosis in Drosophila melanogaster.” Apoptosis 24(11-12): 972–989.

Heisenberg, M. (1998). “What do the mushroom bodies do for the insect brain? an introduction.” Learn Mem 5(1-2): 1–10.

Awasaki, T., Y. Huang, M. B. O’Connor and T. Lee (2011). “Glia instruct developmental neuronal remodeling through TGF-β signaling.” Nature neuroscience 14(7): 821–823.

Hakim, Y., S. P. Yaniv and O. Schuldiner (2014). “Astrocytes Play a Key Role in Drosophila Mushroom Body Axon Pruning.” PLOS ONE 9(1): e86178.

Yu, X. M., I. Gutman, T. J. Mosca, T. Iram, E. Ozkan, K. C. Garcia, L. Luo and O. Schuldiner (2013). “Plum, an immunoglobulin superfamily protein, regulates axon pruning by facilitating TGF-β signaling.” Neuron 78(3): 456–468.

MacDonald, J. M., M. G. Beach, E. Porpiglia, A. E. Sheehan, R. J. Watts and M. R. Freeman (2006). “The Drosophila cell corpse engulfment receptor Draper mediates glial clearance of severed axons.” Neuron 50(6): 869–881.

Doherty, J., M. A. Logan, O. E. Tasdemir and M. R. Freeman (2009). “Ensheathing Glia Function as Phagocytes in the Adult Drosophila Brain.” Journal of Neuroscience 29(15): 4768–4781.

Boulanger, A., C. Thinat, S. Züchner, L. G. Fradkin, H. Lortat-Jacob and J.-M. Dura (2021). “Axonal chemokine-like Orion induces astrocyte infiltration and engulfment during mushroom body neuronal remodeling.” Nature Communications 12(1): 1849.

Ji, H., B. Wang, Y. Shen, D. Labib, J. Lei, X. Chen, M. Sapar, A. Boulanger, J. M. Dura and C. Han (2023). “The Drosophila chemokine-like Orion bridges phosphatidylserine and Draper in phagocytosis of neurons.” Proc Natl Acad Sci U S A 120(24): e2303392120.

Perron, C., P. Carme, A. L. Rosell, E. Minnaert, S. Ruiz-Demoulin, H. Szczkowski, L. J. Neukomm, J.-M. Dura and A. Boulanger (2023). “Chemokine-like Orion is involved in the transformation of glial cells into phagocytes in different developmental neuronal remodeling paradigms.” Development 150(19).

Schafer, D. P., E. K. Lehrman, A. G. Kautzman, R. Koyama, A. R. Mardinly, R. Yamasaki, R. M. Ransohoff, M. E. Greenberg, B. A. Barres and B. Stevens (2012). “Microglia sculpt postnatal neural circuits in an activity and complement-dependent manner.” Neuron 74(4): 691–705.

Yaniv, S. P. and O. Schuldiner (2016). “A fly’s view of neuronal remodeling.” Wiley Interdiscip Rev Dev Biol 5(5): 618–635.

Boulanger, A. and J.-M. Dura (2022). “Neuron-glia crosstalk in neuronal remodeling and degeneration: Neuronal signals inducing glial cell phagocytic transformation in Drosophila.” BioEssays 44(5): 2100254.

Pfeiffer, B. D., A. Jenett, A. S. Hammonds, T.-T. B. Ngo, S. Misra, C. Murphy, A. Scully, J. W. Carlson, K. H. Wan, T. R. Laverty, C. Mungall, R. Svirskas, J. T. Kadonaga, C. Q. Doe, M. B. Eisen, S. E. Celniker and G. M. Rubin (2008). “Tools for neuroanatomy and neurogenetics in *Drosophila*.” Proceedings of the National Academy of Sciences 105(28): 9715–9720.

Jenett, A., Gerald, T.-T. B. Ngo, D. Shepherd, C. Murphy, H. Dionne, Barret, A. Cavallaro, D. Hall, J. Jeter, N. Iyer, D. Fetter Joanna, H. Peng, Eric, Robert, Eugene, Zbigniew, Y. Aso, Gina, A. Enos, P. Hulamm, Shing, H.-H. Li, Todd, F. Long, L. Qu, Sean K. Rokicki, T. Safford, K. Shaw, Julie, A. Sowell, S. Tae, Y. Yu and Christopher (2012). “A GAL4-Driver Line Resource for Drosophila Neurobiology.” Cell Reports 2(4): 991–1001.

Manoukian, A. S. and H. M. Krause (1992). “Concentration-dependent activities of the even-skipped protein in Drosophila embryos.” Genes Dev 6(9): 1740–1751.

Nern, A., B. D. Pfeiffer and G. M. Rubin (2015). “Optimized tools for multicolor stochastic labeling reveal diverse stereotyped cell arrangements in the fly visual system.” Proceedings of the National Academy of Sciences 112(22): E2967–E2976.

Viswanathan, S., M. E. Williams, E. B. Bloss, T. J. Stasevich, C. M. Speer, A. Nern, B. D. Pfeiffer, B. M. Hooks, W.-P. Li, B. P. English, T. Tian, G. L. Henry, J. J. Macklin, R. Patel, C. R. Gerfen, X. Zhuang, Y. Wang, G. M. Rubin and L. L. Looger (2015). “High-performance probes for light and electron microscopy.” Nature methods 12(6): 568–576.

Zhu, S., R. Chen, P. Soba and Y.-N. Jan (2019). “JNK signaling coordinates with ecdysone signaling to promote pruning of *Drosophila* sensory neuron dendrites.” Development 146(8): dev163592.

Cherbas, L., X. Hu, I. Zhimulev, E. Belyaeva and P. Cherbas (2003). “EcR isoforms in Drosophila: testing tissue-specific requirements by targeted blockade and rescue.” Development 130(2): 271–284.

Meltzer, H., E. Marom, I. Alyagor, O. Mayseless, V. Berkun, N. Segal-Gilboa, T. Unger, D. Luginbuhl and O. Schuldiner (2019). “Tissue-specific (ts)CRISPR as an efficient strategy for in vivo screening in Drosophila.” Nature Communications 10(1): 2113.

Yao, T. P., B. M. Forman, Z. Jiang, L. Cherbas, J. D. Chen, M. McKeown, P. Cherbas and R. M. Evans (1993). “Functional ecdysone receptor is the product of EcR and Ultraspiracle genes.” Nature 366(6454): 476–479.

Zheng, X., J. Wang, T. E. Haerry, A. Y. H. Wu, J. Martin, M. B. O’Connor, C.-H. J. Lee and T. Lee (2003). “TGF-β Signaling Activates Steroid Hormone Receptor Expression during Neuronal Remodeling in the Drosophila Brain.” Cell 112(3): 303–315.

Kang, J., Sheehan, De La Torre, Jay, Chiao, Hulegaard, Corty, Baconguis. Zhou, Freeman (2023). “Tweek-dependent formation of ER-PM contact sites enables astrocyte phagocytic function and remodeling of neurons.” bioRxiv: 2023.2011.2006.565932.

Poe, A. R., L. Tang, B. Wang, Y. Li, M. L. Sapar and C. Han (2017). “Dendritic space-filling requires a neuronal type-specific extracellular permissive signal in Drosophila.” Proc Natl Acad Sci U S A 114(38): E8062–e8071.

Winbush, A. and J.C. Weeks “Steroid-triggered, cell-autonomous death of a Drosophila motoneuron during metamorphosis.” (1749-8104 (Electronic)).

Denton, D., B. V. Shravage, R. Simin, E. H. Baehrecke and S. Kumar (2010). “Larval midgut destruction in Drosophila: Not dependent on caspases but suppressed by the loss of autophagy.” Autophagy 6(1): 163–165.

Zirin, J., D. Cheng, N. Dhanyasi, J. Cho, J.-M. Dura, K. VijayRaghavan and N. Perrimon (2013). “Ecdysone signaling at metamorphosis triggers apoptosis of Drosophila abdominal muscles.” Developmental Biology 383(2): 275–284.

Peterson, J. S., M. Barkett and K. McCall (2003). “Stage-specific regulation of caspase activity in drosophila oogenesis.” Developmental Biology 260(1): 113–123.

Clem, R. J., M. Fechheimer and L. K. Miller (1991). “Prevention of Apoptosis by a Baculovirus Gene During Infection of Insect Cells.” Science 254(5036): 1388–1390.

Delorenzi, M. and M. Bienz (1990). “Expression of Abdominal-B homeoproteins in Drosophila embryos.” Development 108(2): 323–329.

Bakshi, A., R. Sipani, N. Ghosh and R. Joshi (2020). “Sequential activation of Notch and Grainyhead gives apoptotic competence to Abdominal-B expressing larval neuroblasts in Drosophila Central nervous system.” PLoS Genet 16(8): e1008976.

Clarembaux-Badell, L., P. Baladrón-de-Juan, H. Gabilondo, I. Rubio-Ferrera, I. Millán, C. Estella, F. S. Valverde-Ortega, I. M. Cobeta, S. Thor and J. Benito-Sipos (2022). “Dachshund acts with Abdominal-B to trigger programmed cell death in the Drosophila central nervous system at the frontiers of Abd-B expression.” Dev Neurobiol 82(6): 495–504.

Monedero Cobeta, I., B. Y. Salmani and S. Thor (2017). “Anterior-Posterior Gradient in Neural Stem and Daughter Cell Proliferation Governed by Spatial and Temporal Hox Control.” Curr Biol 27(8): 1161–1172.

Lai, Y.-W., S.-Y. Chu, J.-Y. Wei, C.-Y. Cheng, J.-C. Li, P.-L. Chen, C.-H. Chen and H.-H. Yu (2016). “Drosophila microRNA-34 Impairs Axon Pruning of Mushroom Body γ Neurons by Downregulating the Expression of Ecdysone Receptor.” Scientific Reports 6(1): 39141.

Krämer, R., N. Wolterhoff, M. Galic and S. Rumpf (2023). “Developmental pruning of sensory neurites by mechanical tearing in Drosophila.” Journal of Cell Biology 222(3).

Arya, R., T. Sarkissian, Y. Tan and K. White (2015). “Neural stem cell progeny regulate stem cell death in a Notch and Hox dependent manner.” Cell Death Differ 22(8): 1378–1387.

Chen, Y.-T., C.-H. Kan, H. Liu, Y.-H. Liu, C.-C. Wu, Y.-P. Kuo, I. Y.-F. Chang, K.-P. Chang, J.-S. Yu and B. C.-M. Tan (2022). “Modular scaffolding by lncRNA HOXA10-AS promotes oral cancer progression.” Cell Death & Disease 13(7): 629.

