## Supplemental Table 1 for "Astrocyte-dependent local neurite pruning and Hox gene-mediated cell death in Beat-Va neurons"

| Line | Gene | Pattern (vs= very simple, s=simple, c=complex, sc=intermediate) | BDSC | Notes |
| --- | --- | --- | --- | --- |
| *GMR52G09-Gal4* | *CG42543 mp* | sc | 38845 |  |
| *GMR55E01-Gal4* | *CG31235* | sc | 39117 |  |
| *GMR58G04-Gal4* | *CG1522 cac* | sc | 39194 |  |
| *GMR64D10-Gal4* | *CG5907 Frq2* | s | 39305 |  |
| *GMR68B07-Gal4* | *CG7485 TyrR* | sc | 39459 | Weak expression |
| *GMR74E08-Gal4* | *CG7847 sr* | s | 39859 |  |
| *GMR75H03-Gal4* | *CG34340* | s | 39908 |  |
| *GMR77E07-Gal4* | *CG34378 Pvf3* | c | 39969 | Very bright |
| *GMR78B07-Gal4* | *CG42281 bun* | sc | 39989 |  |
| *GMR83B05-Gal4* | *CG7994 5-HT2B* | vs | 40352 | No GFP expression |
| *GMR84D10-Gal4* | *CG18389 Eip93F* | sc | 40392 |  |
| *GMR85C10-Gal4* | *CG1343 Sp1* | sc | 40424 | Bright |
| *GMR85F10-Gal4* | *CG7734 shn* | s | 40434 |  |
| *GMR91C05-Gal4* | *CG42242 beat-VII* | sc | 40578 | Weak expression |
| *GMR93G03-Gal4* | *CG31247 tinc* | s | 40660 |  |
| *GMR93G08-Gal4* | *CG42625 mun* | c | 40664 | Bright |
| *GMR94E01-Gal4* | *CG8254 exex* | s | 40683 |  |
| *GMR94G06-Gal4* | *CG33512 dpr4* | sc | 40701 | Bright |
| *GMR41F12-Gal4* | *CG7902 bap* | s | 41242 |  |
| *GMR21F04-Gal4* | *CG14307 fru* | s | 45462 | No GFP expression |
| *GMR32F03-Gal4* | *CG7100 CadN* | s | 45588 |  |
| *GMR13E09-Gal4* | *CG1470 Gycbeta100B* | vs | 45797 | Weak expression |
| *GMR39H05-Gal4* | *CG1832 Clamp* | s | 45873 |  |
| *GMR46H01-Gal4* | *CG5954 l(3)mbt* | s | 45978 |  |
| *GMR18A10-Gal4* | *CG11325 GRHR* | s | 46156 |  |
| *GMR50G02-Gal4* | *CG3114 ewg* | s | 46282 |  |
| *GMR56G03-Gal4* | *CG3143 foxo* | s | 46336 | Weak expression |
| *GMR72D03-Gal4/TM3, Sb* | *CG33517 D2R* | c | 46676 | No GFP expression |
| *GMR82A06-Gal4/TM3, Sb* | *CG11152 fd102C* | c | 47111 |  |
| *GMR95A04-Gal4* | *CG12769* | c | 47271 | Bright |
| *GMR44B11-Gal4* | *CG5133 Doc1* | s | 47358 |  |
| *GMR52C09-Gal4* | *CG6669 klg* | c | 47371 |  |
| *GMR54F07-Gal4* | *CG1522 cac* | s | 47377 | Weak expression |
| *GMR59C02-Gal4* | *CG5744 Frq1* | c | 47383 | Bright |
| *GMR65H09-Gal4* | *CG7887 Takr99D* | sc | 47389 |  |
| *GMR74A06-Gal4* | *CG17390 Oaz* | s | 47398 | Bright |
| *GMR1E307-Gal4* | *CG6494 h* | vs | 47861 | Weak expression |
| *GMR21H03-Gal4* | *CG1916 Wnt2* | s | 47900 |  |
| *GMR36H01-Gal4* | *CG17888 Pdp1* | s | 47920 |  |
| *GMR46C12-Gal4* | *CG17077 pnt* | s | 47938 | Bright |
| *GMR58H02-Gal4* | *CG3967* | c | 47952 | Bright |
| *GMR94A09-Gal4* | *CG5518 sda* | c | 48007 | No GFP expression |
| *GMR24G12-Gal4* | *CG7807 AP-2* | s | 48053 | Nice pattern |
| *GMR29E07-Gal4* | *CG9019 dsf* | s | 48090 |  |
| *GMR29H05-Gal4* | *CG1634 Nr* | sc | 48094 | Bright |
| *GMR31C03-Gal4* | *CG7100 CadN* | c | 48103 | Bright |
| *GMR40G11-Gal4* | *CG33956 kay* | s | 48143 | Weak expression |
| *GMR54E11-Gal4/TM3, Sb* | *CG5744 Frq1* | c | 48203 |  |
| *GMR85G10-Gal4* | *CG7734 shn* | s | 48384 | Weak expression |
| *GMR88D01-Gal4/TM3, Sb* | *CG32447* | s | 48395 | No GFP expression |
| *GMR94A04-Gal4/TM3, Sb* | *CG5518 sda* | vs | 48423 |  |
| *GMR10D05-Gal4* | *CG12287 pdm2* | c | 48438 |  |
| *GMR12G06-Gal4* | *CG32474 dys* | c | 48524 |  |
| *GMR14A08-Gal4* | *CG8095 scb* | s | 48594 | Messy labeling |
| *GMR76C04-Gal4/TM3, Sb* | *CG12506* | c | 48621 | Bright |
| *GMR91E03-Gal4* | *CG34385 dpr12* | s | 48631 |  |
| *GMR92H04-Gal4/TM3, Sb* | *CG10134 beat-Va* | sc | 48632 |  |
| *GMR92H12-Gal4/TM3, Sb* | *CG4846 beat-Ia* | s | 48633 | Weak expression |
| *GMR14G09-Gal4* | *CG10844 Rya-r44* | c | 48662 | Inconsistent |
| *GMR17C08-Gal4* | *CG10037 vvl* | c | 48761 | Bright |
| *GMR18A04-Gal4* | *CG12370* | c | 48793 | Bright |
| *GMR19F05-Gal4* | *CG7665 Fsh* | c | 48855 | Weak expression |
| *GMR20B01-Gal4* | *CG6383 crb* | s | 48877 | Inconsistent |
| *GMR21D12-Gal4* | *CG9554 CG2302 nAcRalpha-7E* | s | 48946 | Bright |
| *GMR23E05-Gal4* | *CG17299 SNF4Agamma* | s | 49029 |  |
| *GMR23F06-Gal4* | *CG30106* | c | 49036 | Inconsistent |
| *GMR25H08-Gal4* | *CG13777 milt* | s | 49146 |  |
| *GMR26H11-Gal4* | *CG9656 grn* | sc | 49206 | Weak expression |
| *GMR21A11-Gal4/TM3, Sb* | *CG9554 eya* | c | 49292 | Bright |
| *GMR33G09-Gal4* | *CG10699 Lim3* | s | 49365 | Messy labeling |
| *GMR50H06-Gal4* | *CG4684 nwk* | c | 49393 |  |
| *GMR60A01-Gal4* | *CG32296 Mrtf* | sc | 49403 |  |
| *GMR70A01-Gal4* | *CG12073 5-HT7* | c | 49414 | Bright |
| *GMR31C11-Gal4* | *CG8355 sli* | s | 49671 | Bright |
| *GMR32G08-Gal4* | *CG18405 Sema-1a* | s | 49729 | Weak expression |
| *GMR33D08-Gal4* | *CG1856 ttk* | c | 49747 | Bright |
| *GMR34E05-Gal4* | *CG10704 toe* | s | 49789 | Bright |
| *GMR19E02-Gal4* | *CG9885 dpp* | c | 49833 | Bright |
| *GMR35F03-Gal4* | *CG1864 Hr38* | sc | 49914 | Bright |
| *GMR38E10-Gal4* | *CG9704 Nrt* | c | 50009 | Bright |
| *GMR40F08-Gal4* | *CG31666 chinmo* | c | 50095 | Bright |
| *GMR40H02-Gal4* | *CG12690 CHES-1-like* | sc | 50102 | Bright |
| *GMR41E11-Gal4* | *CG11020 nompC* | c | 50131 | Bright |
| *GMR42E12-Gal4* | *CG11153 Sox102F* | s | 50159 | Bright |
| *GMR44D10-Gal4* | *CG5685 Calx* | c | 50209 | Bright |
| *GMR45A05-Gal4* | *CG5695 jar* | s | 50218 | Bright |
