## Supplemental Table 2 for "Astrocyte-dependent local neurite pruning and Hox gene-mediated cell death in Beat-Va neurons"

| Line | Gene | Pattern (s=simple, c=complex, sc=intermediate) | BDSC | Notes |
| --- | --- | --- | --- | --- |
| *GMR52G09-Gal4* | *CG42543  mp* | sc | 38845 | Good larval and 6APF expression. Gone by 18APF. |
| *GMR58G04-Gal4* | *CG1522  cac* | sc | 39194 | Faint larval expression, most cells disappear by 6APF. |
| *GMR64D10-Gal4* | *CG5907  Frq2* | s | 39305 | Bright cell bodies at larva, no expression at 6 APF, maybe new cells express GFP by 18 APF |
| *GMR74E08-Gal4* | *CG7847  sr* | s | 39859 | Not good expression. |
| *GMR75H03-Gal4* | *CG34340* | s | 39908 | Strong larval expression. Gone by 6APF. |
| *GMR77E07-Gal4* | *CG34378  Pvf3* | c | 39969 | Good expression throughout metamorphosis. |
| *GMR78B07-Gal4* | *CG42281  bun* | sc | 39989 | Simple pattern, expresses throughout metamorphosis. |
| *GMR85C10-Gal4* | *CG1343  Sp1* | sc | 40424 | Only pictures for wL3wL3 and 6APF. Strong GFP expression. |
| *GMR85F10-Gal4* | *CG7734  shn* | s | 40434 | Probably labels motor neurons. Not a lot of refinement. |
| *GMR93G03-Gal4* | *CG31247  tinc* | s | 40660 | Only pictures for wL3 and 6APF. Labels 4 cells. Expression fades. |
| *GMR94E01-Gal4* | *CG8254  exex* | s | 40683 | Strong expression. Not a lot of refinement. |
| *GMR18A10-Gal4* | *CG11325  GRHR* | s | 46156 | Weak larval expression, strong at 18APF. |
| *GMR50G02-Gal4* | *CG3114  ewg* | s | 46282 | Weak expression. |
| *GMR72D03-Gal4/TM3, Sb* | *CG33517  D2R* | c | 46676 | Good expression throughout metamorphosis. Maybe cell death candidate. |
| *GMR82A06-Gal4/TM3, Sb* | *CG11152 fd102C* | c | 47111 | Labels 6 cells. Not a lot of refinement through 6APF. |
| *GMR95A04-Gal4* | *CG12769* | c | 47271 | Good expression throughout. Maybe cell death candidate. |
| *GMR44B11-Gal4* | *CG5133  Doc1* | s | 47358 | Good expression throughout. Sparse labeling Increases in cell number at 18APF. |
| *GMR52C09-Gal4* | *CG6669  klg* | c | 47371 | Good expression throughout. Cell death and perhaps refinement. |
| *GMR74A06-Gal4* | *CG17390  Oaz* | s | 47398 | Good expression. Not a lot of refinement. |
| *GMR29H05-Gal4* | *CG1634  Nr* | sc | 48094 | Probably labels motor neurons. Not a lot of refinement. |
| *GMR31C03-Gal4* | *CG7100  CadN* | c | 48103 | Probably labels motor neurons. Some refinement. |
| *GMR92H04-Gal4/TM3, Sb* | *CG10134  beat-Va* | sc | 48632 | Good expression at 6APF and wL3. |
| *GMR21D12-Gal4* | *CG9554  CG2302  nAcRalpha-7E* | s | 48946 | Not a lot of refinement, maybe an increase in cell number. |
| *GMR25H08-Gal4* | *CG13777  milt* | s | 49146 | Not a lot of refinement. 4 cells/side. |
| *GMR21A11-Gal4/TM3, Sb* | *CG9554  eya* | c | 49292 | Good expression + refinement. |
| *GMR35F03-Gal4* | *CG1864  Hr38* | sc | 49914 | Messy. |
| *GMR42E12-Gal4* | *CG11153  Sox102F* | s | 50159 | Inconsistent labeling. |
